# Electrophysiological brain reconfiguration during meditation-induced extended cessation of consciousness: an EEG microstate study

**DOI:** 10.64898/2026.02.10.705005

**Authors:** David A.E. Zarka, Winson F.Z. Yang, Abel Rassat, Ruby M. Potash, Terje Sparby, Matthew D. Sacchet

**Author notes:** Corresponding author: Matthew D. Sacchet, Ph.D., **E-mail:**. Denotes co-second authors (Winson F.Z. Yang, Abel Rassat).

## Abstract

*Extended cessation* (EC) is a rare, non-ordinary meditative endpoint characterized by a temporary absence of reportable phenomenal experience, followed by an extraordinary perceptual vividness, openness, equanimity and affective balance. As such, EC provides a unique, non-pharmacological model for investigating the neural basis of the suspension and restoration of consciousness. Here, we examined whole-brain electrophysiological activity during EC using dense-sampling electroencephalographic microstate analysis in five highly trained meditators. Temporal parameters and transition probabilities of canonical microstates during EC were compared with two control conditions (counting and memory tasks) across six frequency bands (broadband, delta, theta, alpha, beta, gamma). EC was associated with a large-scale reconfiguration of brain dynamics, characterized by reduced patterns linked to self-related processing (microstate B) and increased temporal prominence of configurations associated with integrative brain activity (microstate C). Transition dynamics indicated a redistribution of information flow, with sensory-related activity (microstate A) preferentially routed toward integrative processes rather than toward self-related processing. Frequency-specific modulation further supported this reorganization, suggesting a decoupling of self-related dynamics alongside changes in local sensory-sensorimotor processing. Together, these findings identify a distinct neural signature of EC at scalp level and support the view that conscious experience can be transiently suspended through a coordinated reconfiguration of large-scale brain dynamics rather than a breakdown of integrative function. More broadly, this work provides a framework for understanding how advanced meditation can flexibly reshape the organization of brain activity, with implications for theories of consciousness as well as mental health, well-being, and flourishing.

**SIGNIFICANCE STATEMENT:** How can conscious experience be voluntarily suspended without loss of wakefulness or brain function? *Extended cessation*, a rare state achieved through advanced meditation, offers a unique opportunity to address this question without pharmacological manipulation. Using high-density EEG, we identify a reliable large-scale reorganization of brain activity during this state, characterized by a reduction of self-related processing and a redistribution of ongoing dynamics toward sensory and integrative activity. These findings show that conscious experience can be transiently disengaged through coordinated changes in brain dynamics rather than global neural shutdown. More broadly, this work provides a new model for studying flexible control over conscious experience, with potential implications for theories of consciousness and mental health.

## INTRODUCTION

Conventional scientific and clinical frameworks generally treat wakefulness as sufficient for the presence of subjective consciousness (1). However, both clinical neurology and contemplative traditions describe rare states in which wakefulness and arousal are preserved while volitional access to experience is profoundly disrupted (2–4). In clinical contexts, this dissociation is illustrated by conditions such as absence seizures or akinetic mutism, where sensory processing may be preserved while individuals are behaviorally unresponsive or show no overt signs of conscious awareness (4). Reports from advanced meditation traditions also describe rare, non-pathological instantiations of such dissociation, in which perceptual experience is profoundly altered (2, 3), becomes temporarily phenomenologically inaccessible (5–8), and is followed by rapid and reliable recovery. In exceptional cases, known as *extended cessation* (EC), perceptual experience is reported to be absent for extended durations ranging from minutes to several hours (9). Crucially, unlike clinical unresponsive states, arousal, postural control, and physiological stability are preserved during EC, and both entry into and emergence from the state are reported as volitional (10). These reports raise foundational questions about the relationship between brain activity, arousal and conscious experience (11, 12).

While a persistent difficulty in this field concerns how “consciousness” itself is defined and operationalized (1, 13), the present study adopts an operational definition of consciousness as *phenomenal consciousness*, understood as the presence of subjective experience encompassing perceptual content, affective tone, and self-referential awareness. Under this definition, phenomenal consciousness may be temporarily absent even while the organism remains biologically alive and capable of subsequently re-entering conscious experience. Accordingly, this operationalization relies on retrospective reportability and reproducible post-state phenomenology. This distinction between biological wakefulness and phenomenal consciousness is articulated explicitly in early Buddhist psychological frameworks, within a vast contemplative literature and phenomenological taxonomy (14). In these traditions, the meditative attainment known as *nirodha samāpatti* is described as a temporary suspension of all phenomenal experience, including affective tone and perceptual recognition, two mental factors regarded as universally present in conscious cognition (9, 15–17). Upon emergence from this state, consciousness is reported to resume with pronounced clarity, equanimity, and affective balance. Historically, *nirodha samāpatti* has been regarded as one of the highest meditative attainments within certain Theravāda contemplative taxonomies (14, 18, 19). For a detailed historical and theoretical discussion, see Laukkonen et al. (9).

In the present study, we investigate *extended cessation* (EC) as a broader phenomenon without making definitive claims about its doctrinal interpretation. Although historical descriptions of EC are related to *nirodha samāpatti*, our focus is on formulating EC in operational terms that permit neuroscientific investigation, rather than resolving questions of historical interpretation or religious taxonomy (20). We define EC as a state that is voluntarily initiated and temporally bounded, in which phenomenal experience is reported to be absent and followed by spontaneous restoration of consciousness and reliable post-state psychological changes. These aftereffects have been described to include exceptional clarity, deep equanimity, marked reductions in mental suffering and repetitive thought, and a sustained absence of inner verbal narration (9). Importantly, EC is distinguished from pharmacologically induced disruptions of consciousness (e.g. anesthesia or psychedelics) in that it is endogenously generated and does not involve exogenous agents (21, 22). As such, EC is defined here at the level of phenomenal consciousness only, and does not imply an absence of brain activity, arousal, or neural processes supporting later recovery of experience.

From a neuroscientific perspective, EC provides a rare opportunity to study the fundamental dynamics of consciousness. Current approaches to studying consciousness or loss of consciousness such as sleep, anesthesia, or pathological disorders, lack volitional control, physiological stability, or prospective timing (21, 22), making it difficult to isolate mechanisms that dismantle and reinstate awareness (9). These exogenous or pathological approach complicate inferences about the neural basis of conscious experience itself. In contrast, EC can be entered and exited intentionally, with quick and discontinuous in- and out-transition phases, precise control over duration, and without external stimulation (9, 23). As such, EC offers a natural and unique model to investigate the brain dynamics underlying consciousness (5, 9). The present definition, however, does not claim that EC is categorically separable from all other non-experiential conditions, but rather delineates an empirical case for investigating changes in phenomenal consciousness. Additionally, the positive aftereffects of EC may relate to substantial neuroplasticity and reorganization of the brain, potentially informing techniques to enhance mental health and overall wellbeing.

In particular, recent advances in computational neuroscience offer a valuable framework for interpreting both the EC itself and its aftereffects (24, 25). Within predictive processing frameworks, both phenomenal consciousness and mental wellbeing have been linked to the flexible balance between internally generated expectations and incoming sensory signals. Conscious experience is thought to reflect the brain’s current best inferential model of its inputs, whereas excessive reliance on high-level predictions is associated with reduced psychological flexibility and wellbeing (26, 27). From this perspective, advanced meditations may reflect a controlled shift in this balance, whereby higher-level self-related predictions are relaxed while lower-level processing remains active (25). In its most extreme form, such as EC, this shift may extend deeply through the brain functional hierarchy, reaching the lowest levels (24). This redistribution, often formalized as *precision re-weighting* within the brain’s generative model, may transiently prevent the integration of information into a coherent, reportable experience, while simultaneously enabling the post-state emergence of increased clarity, equanimity, and reduced cognitive rigidity (25).

To date, empirical research on EC remains extremely limited, with only two investigations (28, 29) and one separate conceptual pilot dataset (9). A recent 7T fMRI study (29) provided the first high-resolution neuroimaging evidence, revealing widespread reductions in cortical activity and connectivity during EC alongside increased differentiation within sensory cortices, suggesting that consciousness can be volitionally suspended without global neural shutdown. Complementing this work, a multimodal electroencephalographic-magnetoencephalographic (EEG-MEG) study (28) reported that EC markedly reduced alpha power while paradoxically increasing neural complexity, indicating that the brain remains dynamically organized even in the absence of conscious experience. Together, these pioneering studies highlight EC as a distinct, endogenously induced, and physiologically stable state that challenges current models of unconsciousness.

The current study extends electrophysiological investigation of EC using high-density EEG to highlight large-scale brain dynamics underlying meditation-induced volitional and prolonged cessations of consciousness. In this respect, EEG captures rapid brain-state transitions through stable scalp topography known as *microstates* (MS), which remain briefly stable (around 40-120 ms) before rapidly switching into a new one, even at rest and in the absence of external afferent volleys (30). These discrete, stable EEG topographical patterns are limited in number and identified from data-driven clustering methods applied to the time series of EEG voltage maps (31). These patterns are relatively homogeneous across studies/conditions and typically fall into four to seven canonical patterns (generally labelled A to G) which are thought to encompass the hubs of resting-state functional networks. Temporal fluctuations in microstates are thus thought to indicate millisecond-scale modifications of how the brain orchestrate its activity, providing a window into higher-order integration processes of the brain (31). In this context, microstate analyses have been applied to a broad range of conscious and unconscious conditions, such as mind wandering (32–34), sleep (35, 36), anesthesia (37), or meditation-related states (38–41).

Previous studies suggested that unconscious states such as propofol-induced anesthesia and deep sleep, show an overall increased duration of microstates, lower occurrence, and reduced complexity, reflecting diminished large-scale integration (35–37). In contrast, meditation studies suggested that daily meditation training induces a decrease in microstate durations which was associated with positive psychological change, such as daily increase in felt attentiveness and serenity (40, 41). As such, decreased microstate duration has been interpreted as reflecting increased lability of the brain networks affording cortical functions greater flexibility and moment-to-moment adaptability (40). Microstate C, associated with the default mode network (DMN) (42) and self-referential internal mentation (43), appears to be particularly sensitive to meditation (41, 44). However, contrasting data suggest that various meditation types differentially affect the temporal parameters of this microstate (38, 39). Notably, transcendental meditation has been shown to decrease the occurrence of microstate C (39), while deep focus attention meditation increases its occurrence and duration (38).

According to previous studies, we assumed that EC relies on a hierarchical reconfiguration of integration processes across functionally segregated areas underlying the conscious activity of the brain and resulting in transformative aftereffects (28, 29). As such, EC should be associated with a radical reorganization of large-scale networks, that should be reflected at the scalp level by altered temporal dynamics of microstates. We hypothesized that EC should lead to an overall decrease in microstate duration and to changes in transition probabilities, especially those related to self-processing and meditation practices such as microstate C. As EC is characterized by a reduced occipital alpha power and increased complexity (28), we expected these changes to occur primarily in the alpha frequency band although not exclusively. This study aims to provide important evidence on the neuroscience of EC, related fundamental aspects of consciousness, and their potential implications for human mental health, resilience and wellbeing.

## RESULTS

### Overview of EEG signals

Figure 1 illustrates near-raw EEG data, including representative EEG traces, time–frequency dynamics, and large-scale functional indices across conditions. As shown in Figure 1A, preprocessed EEG signals exhibit readily distinguishable patterns between conditions, with EC characterized by reduced rhythmic activity compared to Memory and Counting conditions. Consistent with this observation, time–frequency representations across participants (Fig. 1B) reveal a marked reduction in alpha-band power during EC, while activity in non-alpha frequencies remains comparatively preserved. Despite these spectral changes, fronto-parietal coherence profiles appear largely comparable across conditions (Fig. 1C), suggesting that large-scale functional coupling is not globally disrupted during EC. Together, these observations situate EC within a distinct electrophysiological regime characterized by attenuated alpha power without widespread breakdown of large-scale coordination, consistent with previous findings (28). In the following sections, we build on this descriptive level by quantifying these large-scale dynamics using EEG microstate analysis across both broadband and narrow frequency bands (Fig. 2-5). For clarity, all results are reported here for the comparison between EC and the combined control condition (CC), while detailed results separating the two control conditions are provided in the Supporting Information Appendix (Suppl. Fig. C-E and Table C-H) to further characterize condition-specific microstate dynamics.

**Figure 1.**
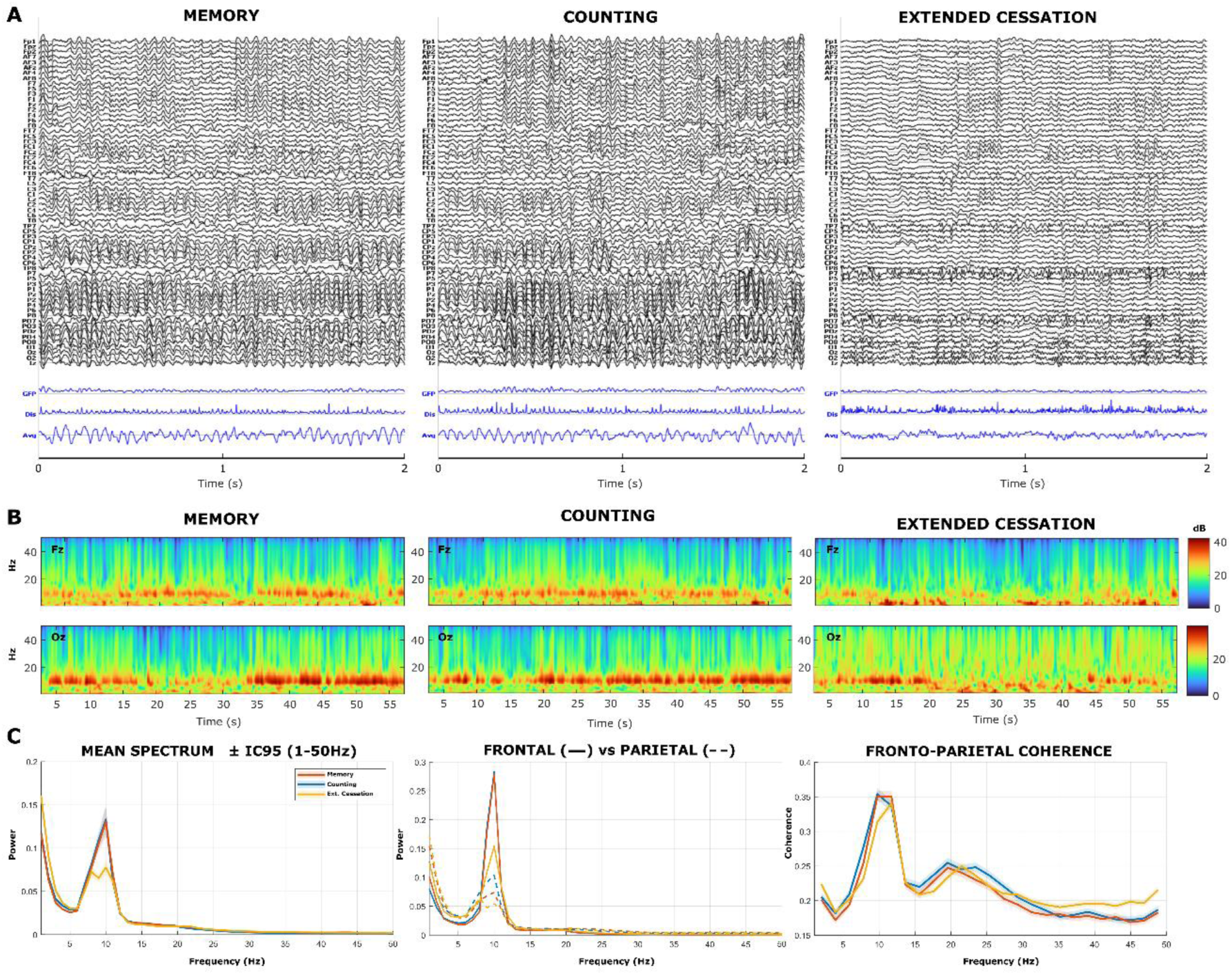
Electrophysiological overview of EEG activity across conditions. **(A)** Representative 2-sec segments of preprocessed EEG recorded during the Memory, Counting, and Extended Cessation (EC) conditions. Global field power (GFP), dissimilarity (Diss), and mean signal (Avg) are displayed below each trace. EC shows reduced oscillatory structure compared to control tasks. **(B)** Time–frequency representations at frontal (Fz) and occipital (Oz) electrodes across conditions. Spectral power (dB) reveals marked attenuation of alpha-band activity (∼8–12 Hz) during EC relative to controls. **(C)** Group-level spectral summaries across conditions. Left: mean power spectrum (1–50 Hz, 95% confidence intervals). Middle: frontal (solid lines) and parietal (dashed lines) power spectra. Right: fronto-parietal coherence. EC is characterized by reduced alpha power while large-scale fronto-parietal coherence remains largely preserved.

**Figure 2.**
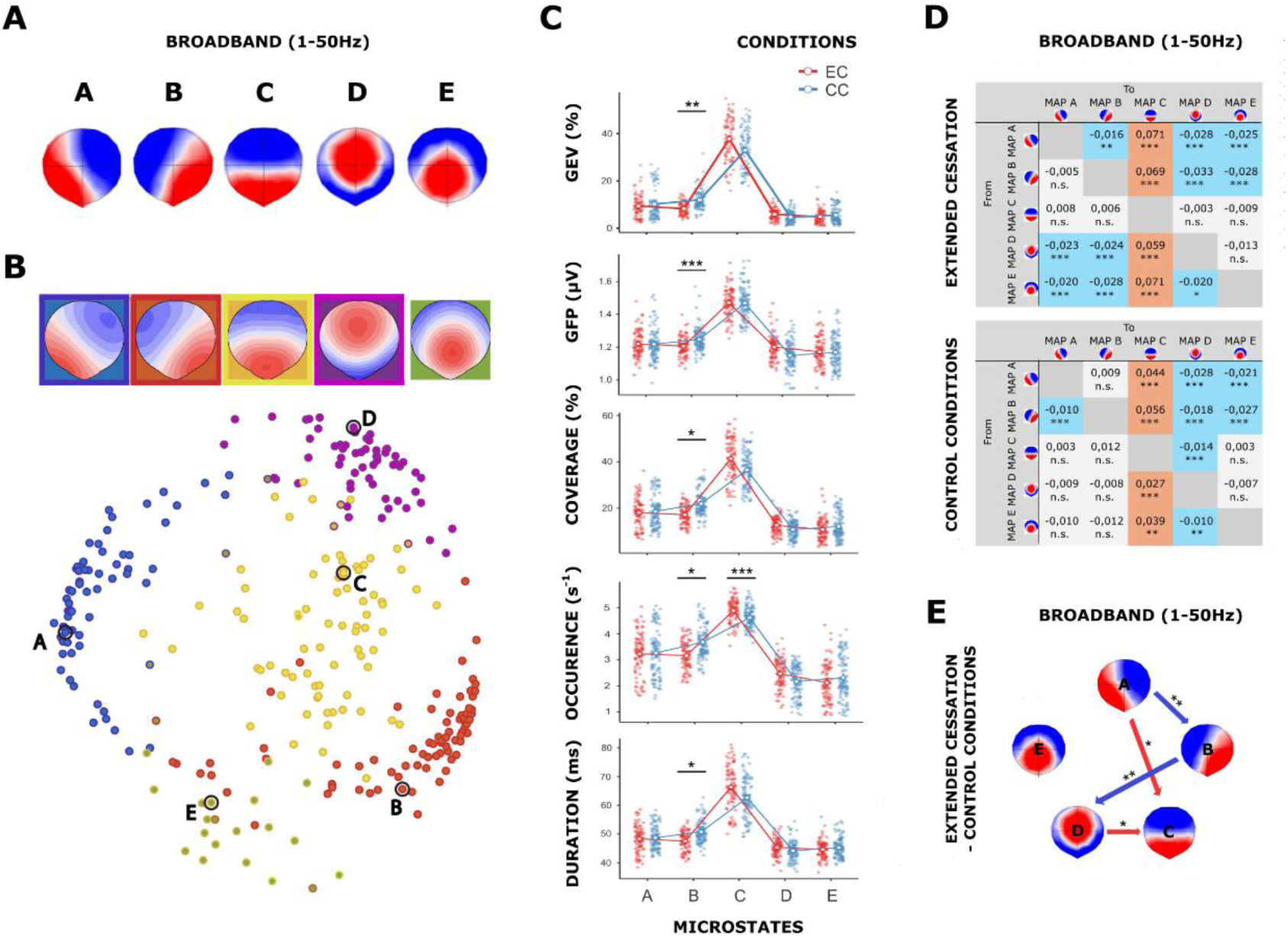
Broadband microstate organization and dynamics (five-microstate solution). **(A)** Group-level broadband microstate topographies (A–E) identified by k-means clustering across participants and conditions. The five canonical maps correspond to established microstate classes characterized by diagonal (A, B), anterior–posterior (C), and centrally symmetric topographies (D, E). **(B)** Meta-microstate validation of the four- and five-microstate solutions. Top: meta-microstate templates for the four-(outlined) and five-microstate (filled) solutions. Bottom: multidimensional scaling of the spatial similarity matrix relating database templates and the current results. Filled and outlined markers represent the five- and four-microstate solutions, respectively. Notably, the four-microstate solution shows that microstate C largely overlaps with microstate E, suggesting a merging of these two topographies under reduced model dimensionality. **(C)** Microstate parameters for extended cessation [EC] and combined control conditions [CC]. EC is characterized primarily by reduced MS B and increased MS C contributions. **(D)** Observed-minus-expected Markov transition probabilities for broadband data (1–50 Hz) in EC (top) and CC (bottom). Warm and cool colors indicate transitions occurring more or less frequently than expected by chance, respectively. **(E)** Significant between-condition differences in transition probabilities. Arrows indicate increased (red) or decreased (blue) transitions during EC. EC was associated with reduced transitions toward MS B and increased transitions toward MS C. Transitions involving microstate E were not significantly altered, suggesting a limited contribution of this microstate to transition reorganization during EC.

### Broadband microstate organization across conditions

Figure 2 illustrates the dominant broadband microstate topographies identified by the *k*-means clustering across all participants and conditions. The optimization metacriterion determined that the optimal number of clusters was *k* = 5, explaining 80.85% of the global variance. As shown in Figure 2A, the segmentation yielded the five canonical maps consistent with previous literature (31, 43, 45) and meta-microstate templates (46). These maps were classified according to the established convention into the following categories: left-right and right-left diagonal orientations (A and B), an anterior-posterior orientation (C), and fronto-central and occipito-central maxima (D and E) (31, 47). Meta-microstate analysis of this broadband template (Fig. 2B) confirmed high-shared spatial variance across all identified microstate classes, both within the global database and within the “Consciousness and its Disorders” research branch (Suppl. Table A). When compared with participant- and condition-specific microstate segmentations, one-to-one spatial correlations exceed r *≥* 0.90 topographic similarity for Counting, Memory and participants S1, S3 and S5, whereas lower correspondence was observed for EC as well as S2 and S4 (Suppl. Fig. B). Consistent with this pattern, the optimization metacriterion identified a four-map solution (*k* = 4) as optimal for EC alone, explaining 78.46% of the global variance, indicating that a reduced microstate configuration more closely reflects the intrinsic structure of EC data (Fig. 2B).

### Broadband microstate dynamics and transitions

Regarding global broadband microstate parameters, the omnibus Wald χ^2^ test for the five-microstates solution revealed significant microstate and interaction effects on GEV, coverage, and occurrence (microstate: all *χ^2^*(*3*) *≥ 225.76, p < 0.001;* interaction: all *χ^2^*(*6*) *≥ 96.68, p < 0.001;* Suppl. Table C). Significant microstate effects were also observed for GFP and duration (all *χ^2^*(*3*) *≥ 3,183.02, p < 0.001*). As illustrated in Figure 2C, MS B showed consistently lower GEV and coverage (all *p < 0.018*, *OR* ≤ *0.73*) as well as lower GFP, occurrence and duration (all *p < 0.049*, *GMR* ≤ *0.97)* during EC compared to CC. In contrast, MS C exhibited increased occurrence during EC (*p < 0.001, GMR = 1.06*). Regarding microstate transitions, most transition probabilities between microstates differed significantly from those expected under random transitions in both EC and CC (Fig. 2D). Overall, transition patterns were largely similar across conditions: MS A, B and D transitioned more frequently toward MS C than expected by chance (all *p* < 0.01, *md ≥* 0.027), whereas other significant transitions occurred less frequently than expected (all *p* < 0.05, *md* < - 0.010). When directly comparing EC and CC (Fig. 2E), differences primarily involved transitions toward microstates B and C. Specifically, EC showed reduced A-to-B and B-to-D transition probabilities (A-to-B*: p* = 0.004, *md* = - 0.024; B-to-D*: p* = 0.003, *md* = - 0.015). In contrast, higher transition probabilities from A and D to C were observed during EC (A-to-C*: p = 0.041, md = 0.028;* D-to-C*: p = 0.035, md = 0.037*).

### EC-aligned microstate structure across frequency bands

Given that condition-specific analyses indicated a four-microstate solution as optimal for EC, frequency-specific group-level maps were spatially correlated with the EC-derived centroids to identify prototypical topographies that best captured EC effects across frequency bands and conditions. Notably, the clustering optimization metacriterion consistently identified optimal solutions ranging from *k* = 4 to *k* = 5 across all narrow frequency bands and conditions. As illustrated in Figure 3, the four-microstate solution (*k* = 4) revealed highly consistent topographies across EEG frequency bands, participants, and conditions, each explaining more than 70% of the global variance (mean: 79.14%, SD: 1.40). In particular, the optimization metacriterion identified *k* = 4 as optimal for the alpha band, explaining 81.70% of the global variance. Across segmentations, the four canonical microstate maps (A–D) were consistently recovered, in agreement with both the five-map solution and the broader microstate literature (31, 43, 45).

**Figure 3.**
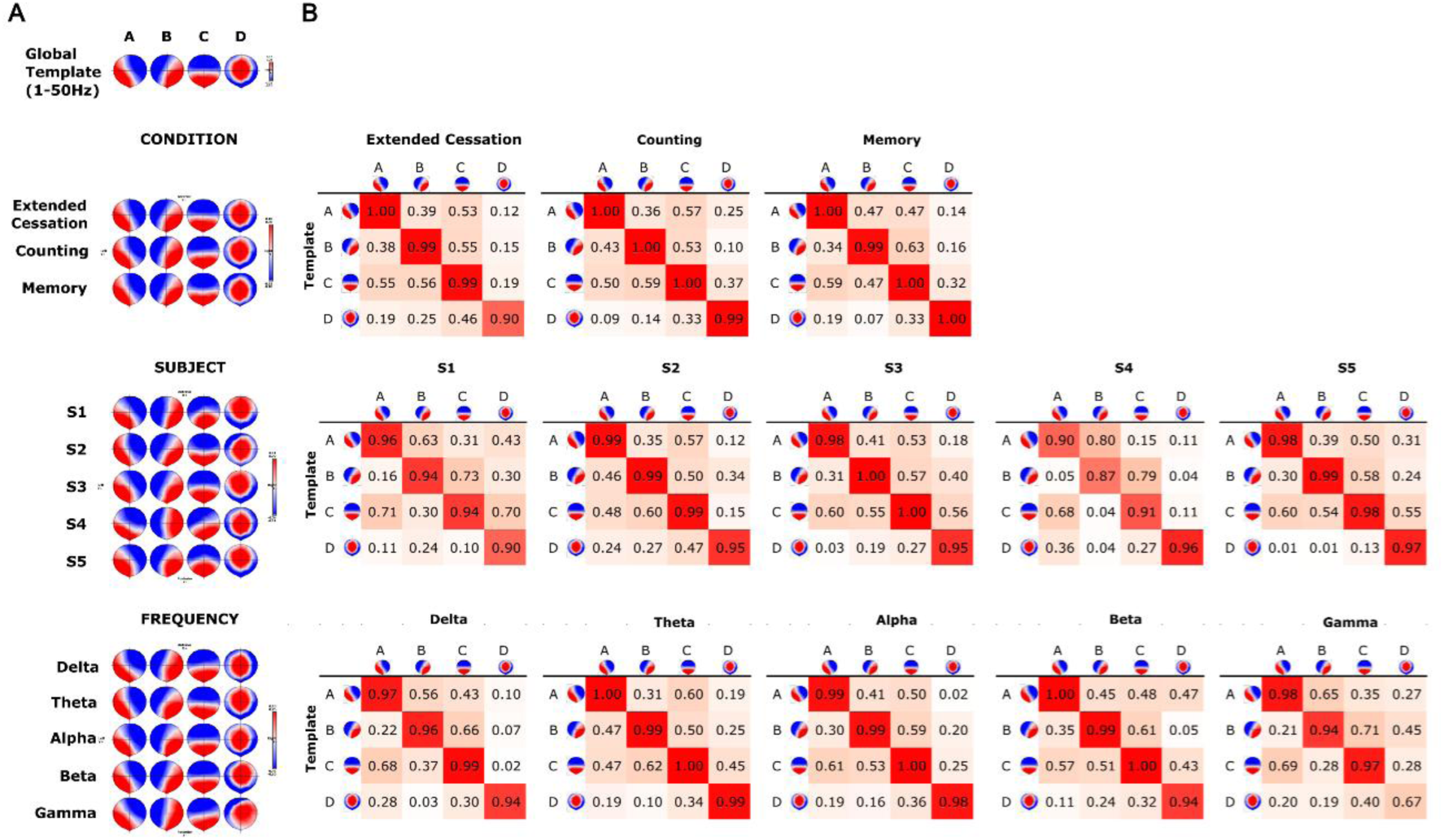
Consistency of four-microstate solution across frequency bands, participants and conditions. **(A)** Canonical broadband microstate topographies (1–50 Hz) and their corresponding maps across experimental conditions (extended cessation, counting, and memory), participants (S1–S5), and frequency bands (delta, theta, alpha, beta, gamma). Spatial configurations remained highly consistent, indicating that EC modulated canonical microstate dynamics rather than inducing novel topographies. **(B)** Spatial correlation matrices between template maps and condition-, subject-, and frequency-specific solutions. High diagonal correlations (highlighted in red) Indicate strong correspondence between matching configurations and confirm the robustness and reproducibility of canonical microstate topographies. These findings support a parsimonious four-microstate solution and suggest that EC primarily alters microstate temporal dynamics and transition structure rather than spatial organization.

When comparing the four-map broadband template with the microstate segmentation obtained for each frequency band, subject and condition, spatial correlations exceeded r *≥* 0.90 topographic similarity in each case (Fig. 3), except for the gamma-MS D (r = 0.67). Meta-microstate analysis further confirmed strong shared spatial variances for all four-map broadband microstate classes, both with the global database and with the “Consciousness and its Disorders” research branch (Suppl. Table B). Notably, the four-map solution indicated that the microstate class corresponding to C exhibited spatial features consistent with both C- and E-like topographies observed in five-map solutions (Fig. 2B), suggesting a partial convergence of these configurations under reduced model dimensionality. Taken together, these results supported the use of a four-microstate solution as a stable and coherent reference across conditions and frequency bands. Consequently, we fitted the four-map broadband templates across all frequency bands to provide a common reference framework for comparing microstate spatio-temporal parameters between conditions in line with established approaches in the field (48). This strategy enabled reliable cross-condition comparisons while accounting for the intrinsic structure of EC.

### Frequency-specific modulation of microstate dynamics

Wald χ^2^ tests revealed significant microstate and interaction effects of amplitude-related (GEV, GFP) and temporal (coverage, occurrence, duration) microstate parameters, indicating robust differences across conditions and frequency bands (all *p* < 0.001; Fig. 4, Suppl. Table D-H). At the broadband level, EC was characterized by a consistent reduction in MS B across all measures, including GEV (*p* = 0.002, OR = 0.71), GFP (*p* = 0.029, GMR = 0.97), coverage (*p* < 0.001, OR = 0.77), occurrence (*p* = 0.002, GMR = 0.90), and duration (*p* = 0.008, GMR = 0.93). In contrast, MS C showed increased GEV (*p* < 0.001, OR = 1.37), coverage (*p* < 0.001, OR = 1.38), occurrence (*p* < 0.001, GMR = 1.04), and duration (*p* < 0.001, GMR = 1.12), consistent with the overall B/C modulation observed in the five-microstate solution.

**Figure 4.**
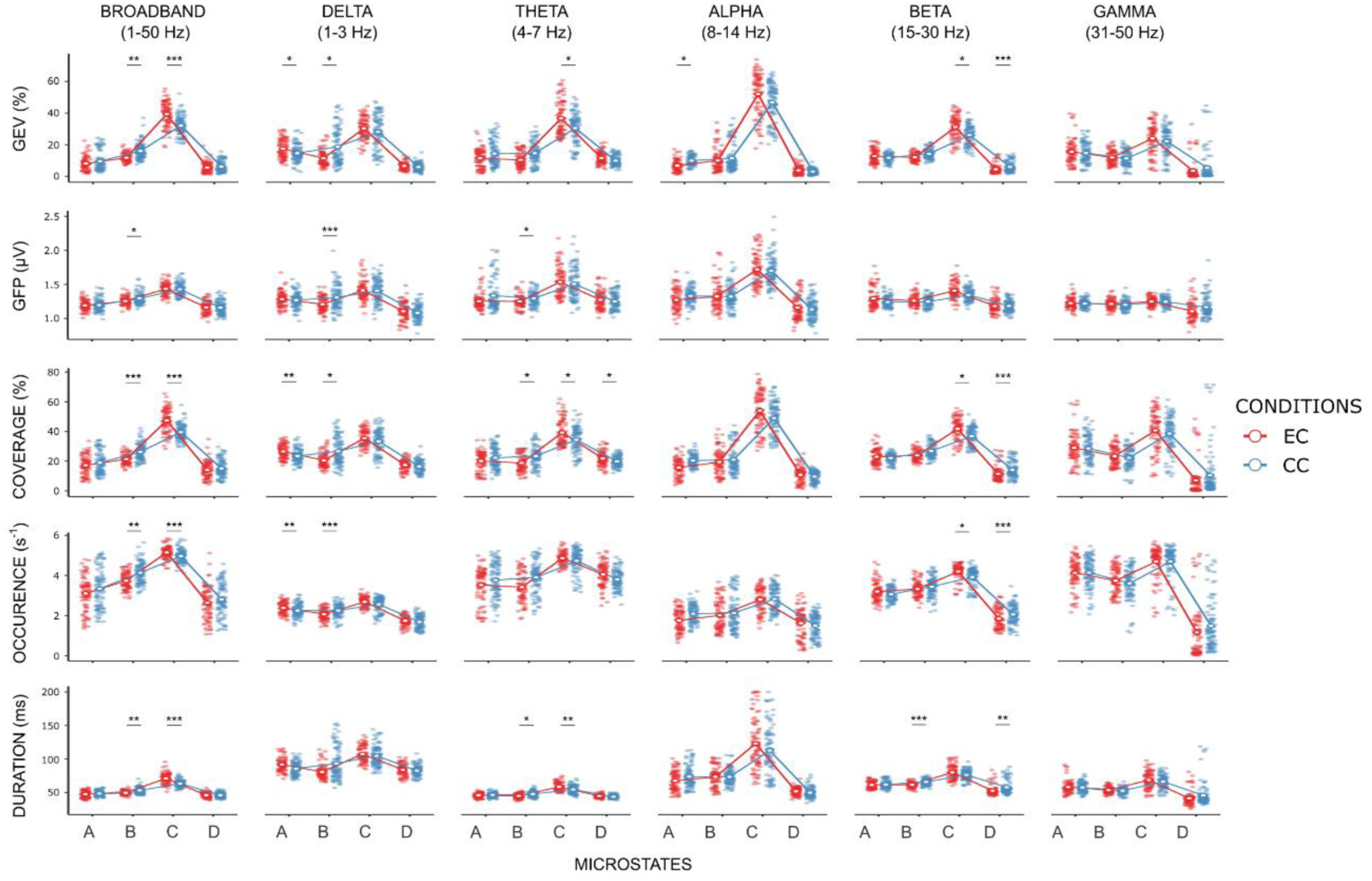
Frequency-specific modulation of microstates parameters during extended cessation. Canonical microstates (A-D) characteristics are shown for broadband (1–50 Hz) and frequency-specific bands (delta, theta, alpha, beta, gamma), comparing extended cessation (EC, red) with the combined control conditions (CC). Rows correspond to global explained variance (GEV), global field power (GFP), time coverage, occurrence rate and mean duration. Statistics were derived from generalized estimating equation (GEE) models with an AR(1) correlation structure and assessed using Wald χ^2^ tests. Asterisks indicate FDR-corrected significance level (* *p* < 0.05, ** *p* < 0.01, *** *p* < 0.001). Results indicate frequency-specific reorganization of microstate dynamics during EC, supporting a model of distributed reconfiguration rather than uniform modulation across oscillatory regimes.

Frequency-specific analyses indicated that these effects were differentially distributed across bands. The reduction in MS B was primarily driven by low-frequency components, with reduced GEV (delta: *p* = 0.010, OR = 0.63), GFP (delta: *p* < 0.001, GMR = 0.93; theta: *p* = 0.036, GMR = 0.97), coverage (delta: *p* = 0.018, OR = 0.75; theta: *p* = 0.036, OR = 0.75), occurrence (delta: *p* < 0.001, GMR = 0.92), and duration (theta: *p* = 0.010, GMR = 0.94; beta: *p* < 0.001, GMR = 0.94) predominantly in delta and theta ranges. Conversely, the increase in MS C was mainly supported by theta and beta bands, with higher GEV (theta: *p* = 0.024, OR = 1.35; beta: *p* = 0.013, OR = 1.28), coverage (theta: *p* = 0.036, OR = 1.24; beta: *p* = 0.015, OR = 1.23), occurrence (beta: *p* = 0.025, GMR = 1.07), and duration (theta: *p* = 0.008, GMR = 1.07). Additionally, more circumscribed effects were observed for other microstates. MS A showed increased contribution in the delta band (GEV: *p* = 0.031, OR = 1.26; coverage: *p* = 0.004, OR = 1.19; occurrence: *p* = 0.005, GMR = 1.06), while MS D exhibited a pattern of increased theta-band coverage (*p* = 0.045, OR = 1.12) but reduced beta-band contribution across GEV (*p* < 0.001, OR = 0.72), coverage (*p* < 0.001, OR = 0.77), occurrence (*p* < 0.001, GMR = 0.89), and duration (*p* = 0.008, GMR = 0.92).

### Frequency-specific modulation of microstate transitions

As illustrated in Figure 5, most transition probabilities between broadband microstate differed significantly from those expected under random transitions in both EC and CC. Overall, transition patterns were largely similar across EC and CC, with MS A, B and D transitioning more frequently toward MS C than expected by chance (all *p* < 0.001, *md* > 0.034). In contrast, all other significant transitions occurred less frequently than expected (except for C-B transitions in EC being more frequent; *p <* 0.010, *md =* 0.018), consistent with the broadband transition pattern observed in the five-microstate solution. A similar overall pattern was observed across narrow frequency bands, with a consistent bias toward transitions to MS C (Fig. 5A). However, some band-specific deviations were present. In the delta-band, EC was associated with increased B-to-A transitions (*p* < 0.001*, md* = 0.060) and decreased C-to-A transitions (*p* = 0.13*, md* = - 0.023) whereas CC showed increased A-to-B transitions (*p* < 0.001*, md* = 0.023). In the theta band, EC exhibited increased C-to-D transitions (*p* = 0.006*, md* = 0.026). In the gamma band, C-to-A transitions were more frequent than expected in both EC (*p* = 0.016*, md* = 0.027) and CC (*p* < 0.001*, md* = 0.027).

Direct comparisons between EC and CC revealed condition-specific differences primarily involving transitions toward microstates C and B (Figure 5B). Specifically, EC showed increased transitions from A and B to C (A-to-C: *p* < 0.001, *md* = 0.044; B-to-C: *p* = 0.040, *md* = 0.029) and decreased A-to-B transitions (*p* = 0.001, *md* = −0.036). Frequency-specific contrasts were more restricted, with significant effects observed in the delta and beta bands (Fig. 5). In the delta band, EC showed increased B-to-A transitions (*p* = 0.032, *md* = 0.039) and decreased transitions involving B and C pathways (D-to-B, B-to-C, C-to-A; all *p* = 0.032, *md* ≤ −0.021). In the beta band, EC was characterized by increased B-to-C transitions (*p* = 0.027, *md* = 0.030) and reduced bidirectional transitions between B and D (all *p* ≤ 0.009, *md* ≤ −0.013).

**Figure 5.**
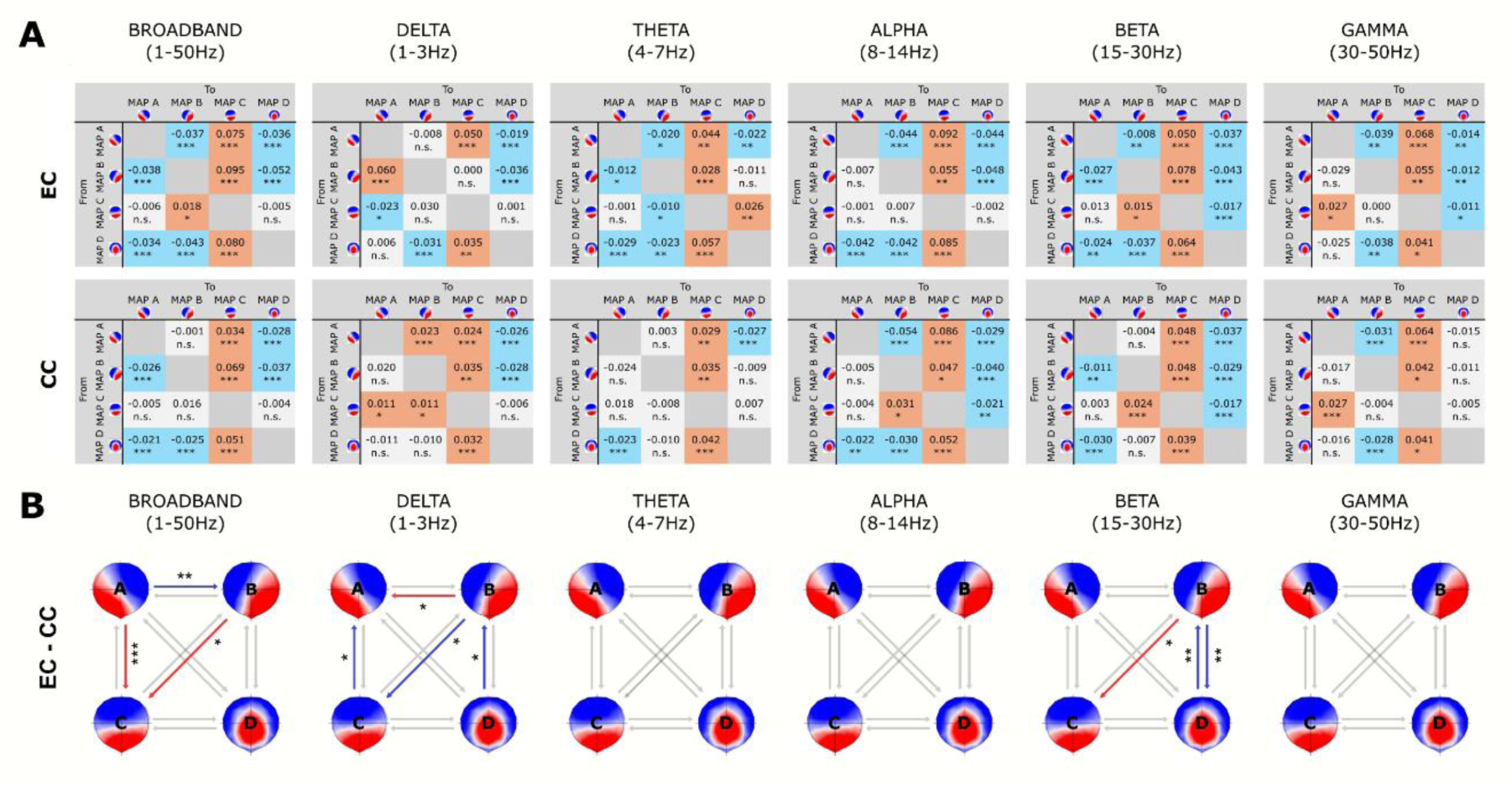
Reconfiguration of microstate transition dynamics during extended cessation. **(A)** Transition probabilities differences (observed-minus-expected) within each condition (Extended Cessation [EC] vs Control Conditions [CC]) across broadband (1-50Hz), delta (1-3Hz), theta (4-7Hz), alpha (8-14Hz), Beta (15-30Hz) and Gamma (30-50Hz) frequency bands. Matrices show transition likelihoods between canonical microstates (rows: “from”, columns: “to”). Positive values (red) indicate more frequent-than-random transitions, whereas negative values (blue) indicate less frequent-than-random transitions. **(B)** Network graphs of between-condition comparisons (EC - CC) in transition probabilities. Arrow indicates transition significantly increased (red) or decreased (blue) during EC relative to CC. Significant effects are indicated by color and text following FDR-adjusted Wald χ^2^ tests (* *p* < 0.05, ** *p* < 0.01, *** *p* < 0.001). The observed transition patterns support a model of directed reorganization of information flow, consistent with altered gating and routing mechanisms across distributed cortical networks.

## DISCUSSION

The present study probes the electrophysiological signatures of an advanced meditative endpoint (14), namely the *extended cessation* of consciousness, using EEG microstate analysis. As expected, EC selectively altered, relative to control conditions, the temporal dynamics of canonical microstates, indicating a shift in the global orchestration of large-scale functional brain networks. In particular, the relative contribution of the microstate labeled as B decreased, whereas that of microstate C increased during EC compared to CC. More specifically, EC was marked by less frequent and shorter occurrences of microstate B, and more frequent occurrences of microstate C. Transition probabilities also reconfigured during EC: transitions from A to C increased, whereas those from A to B decreased. These effects were observed in the broadband EEG, but were distributed across delta, theta, and beta frequency sub-bands, while no significant changes were observed in the alpha band. Additional frequency-specific changes were also observed for microstates A and D. The delta band showed increased occurrence of microstate A and higher B-to-A transitions probabilities during EC, while the beta band showed less frequent and shorter occurrences of microstate D and decreased bidirectional B-to-D transitions. These findings indicate that extended cessation of consciousness expresses a distinctive electrophysiological profile at the scalp and offers a window into the large-scale neural mechanisms that regulate the integration (or suspension) of conscious experience.

EC is distinctive both in its mode of induction and its aftereffects, suggesting that it relies on a specific pattern of functional dissociation. Unlike anesthesia or sleep, which are typically characterized by externally imposed or physiologically driven reductions in consciousness, EC is intentionally induced through extensive practice and can be exited abruptly without residual confusion or drowsiness (9). Instead, practitioners report heightened perceptual clarity, relief, and equanimity, alongside reduced self-referential mental activity upon re-emergence (49). In this respect, EC shares functional similarities with clinical conditions in which conscious experience or volitional engagement is suspended without global neural shutdown, such as absence seizures or akinetic mutism (4). In such conditions, sensory processing may remain preserved yet become decoupled from behavioral integration, and they have been conceptualized as involving selective gating or functional disconnection of integrative systems rather than generalized cortical suppression (4). Within this perspective, the post-EC afterglow of perceptual vividness and clarity may be understood as reflecting forms of cortical hyperexcitability and sensory release observed following transient functional suppression in intact cortex, such as in Charles Bonnet syndrome or post-ictal phenomena following occipital seizures (50, 51). In EC, however, this effect appears tightly constrained and embedded within a controlled re-emergence of conscious awareness.

Within this broader framework, EC may be understood as a rare, non-pathological instance of such selective gating, distinguished by its volitional, reversible, and physiologically stable nature. The combination of reduced alpha power and paradoxically increased neural complexity (28) suggests that EC relies on an extreme form of large-scale alpha gating, potentially impairing the global integration of information into conscious experience while preserving dynamical organization of neuronal activity. Rather than reflecting degraded processing, the transient suspension of phenomenal consciousness during EC may reflect a controlled disengagement or reconfiguration of large-scale integrative networks, while unimodal sensory systems remain locally active and dynamically intact (29). In this view, such gating does not arise from dysfunction, but from the sustained occupation of the gating system by a stable, volitionally maintained top-down control state, which selectively prevents the integration of salient sensory or interoceptive signals into conscious awareness. Within a predictive processing framework, this may reflect a transient, top-down attenuation of high-level, self-referential priors during EC, allowing sensory signals to exert a greater influence on hierarchical reorganization, thereby contributing to the reported enhancement in perceptual clarity and vividness upon re-emergence (25).

Consistent with this large-scale reconfiguration of integrative dynamics, EC exhibits distinctive alterations in the temporal organization of EEG microstates, particularly involving microstate B and C. Specifically, EC was characterized by reduced contribution of microstate B, a class most consistently associated with visual processing involving self-referential components. Converging evidence across neuroimaging and behavioral studies has linked microstate B to internally oriented visual cognition, including self-visualization and autobiographical memory processes (43). Experimental findings further indicate that memory retrieval selectively increases the occurrence of microstate B and enhances C-to-B transition probabilities relative to arithmetic tasks and rest, reflecting a shift toward internally generated self-related imagery (52). In line with this interpretation, clinical populations characterized by altered self-referential processing, such as euthymic bipolar disorder, show reduced occurrence of microstate B alongside increased B-to-C transition probabilities, suggesting an unstable engagement of self-related visual imagery networks (53). Our results closely mirror this pattern: microstate B temporal parameters were significantly reduced during EC, accompanied by increased B-to-C transition probabilities compared to the memory condition (Suppl. Fig. D, E). Taken together, this profile provides converging support for the interpretation that microstate B indexes self-directed autobiographical and imagery-related processes. Accordingly, the reduction of microstate B, coupled with its increased transitions toward microstate C, is consistent with a sustained down-regulation of self-referential processing during EC, aligning with first-person reports of diminished self-related mentation and experiential relief following the cessation state.

Complementing the sustained down-regulation of microstate B, EC was also characterized by a marked increase in the dominance of microstate C, pointing to a coordinated reconfiguration of large-scale functional dynamics. This pattern differs from several meditation studies reporting reduced prevalence of microstate C (39, 54), suggesting that EC engages partially distinct large-scale dynamics within the broader spectrum of meditative states. The increased dominance of microstate C during EC can be interpreted in light of two partially overlapping lines of evidence. First, prolonged dominance of C-like topographies has been associated with reduced responsiveness and executive disengagement in altered states of consciousness. For instance, loss of consciousness under propofol is accompanied by longer microstate durations and reduced sequence complexity, reflecting a breakdown of flexible large-scale coordination (37). A comparable pattern emerges during NREM sleep with dream reports, where increased prevalence of C-like microstate has been interpreted as reflecting reduced executive control alongside preserved internally structured experience (35, 36).

Second, microstate C has been consistently linked to the DMN and internally oriented cognition, including self-oriented mentation (43). In task-based paradigms with experience sampling, episodes of off-task thought are associated with increased coverage (and GEV) of microstate C, accompanied by reciprocal decreases of microstate E, associated with the salience (mid-cingulo-insular) network that supports tonic alertness and task-directed control (55). This dynamic antagonism between microstate C (off-task) and E (on-task) has been demonstrated at the millisecond scale in scalp EEG and is thought to reflect rapid switching between competing functional networks (55). Our findings place EC at an intermediate and functionally distinctive point within this spectrum. The increased occurrence and duration of C (along with increased transition from A and B to C) likely reflect (i) a state of profound inward absorption, in which executive control processes are partially disengaged while internally coherent dynamics are maintained, and (ii) a metastable re-weighting toward DMN-like generators during the deliberate suspension of phenomenal consciousness. This interpretation is consistent with data from meditative absorption in adept practitioners, where microstate C increases, alongside reduced posterior cingulate cortex connectivity and enhanced frontal-temporal coupling, suggesting a decoupling of canonical DMN hubs while preserving distributed coordination (38).

In this context, the apparent convergence between C- and E-like topographies in our data (Fig. 2) provides further insight into this reconfiguration, potentially reflecting the distinctive functional organization of EC. While salience network has been associated with the detection of behaviorally relevant events and the coordination of large-scale network switching, EC may involve the co-expression of salience-related and DMN-like processes within a unified functional regime sustained by strong top-down control, rather than a simple reflecting shift toward internally oriented processing. Such functional overlaps can give rise to highly correlated scalp topographies, making them less separable at the clustering level. This interpretation is consistent with previous findings showing that, under reduced model dimensionality (e.g., *k* = 4), microstates C and E tend to collapse into a single centroid due to their high spatial similarity (42). Importantly, our data suggests that this convergence is not merely a consequence of model reduction, but reflects an underlying reorganization of large-scale networks during EC. Such a reduction in effective dimensionality is consistent with broader observations in altered states of consciousness, where large-scale brain dynamics reorganize toward a more restricted and stabilized set of dominant configurations (56). This view is further supported by evidence from meditation training studies showing reconfigurations in microstate network organization, including altered interactions and integration between large-scale functional systems following contemplative practice (57). Accordingly, the observed dominance of microstate C is best understood not as the expression of a single network, but as the composite expression of salience engagement and DMN re-weighting within a reorganized large-scale functional landscape.

Extending this large-scale reconfiguration, additional EC-specific changes were observed in transition probabilities involving microstate A. Specifically, EC was associated with reduced A-to-B transition probabilities, and increased A-to-C transitions. While still debated, microstate A is generally linked to sensory processing and arousal-related functions (43). A reduction in A-to-B transitions may thus indicate a weakening of the coupling between sensory inflow and networks supporting self-referential and imagery-related processing. In contrast, increased A-to-C transitions suggests a stronger contribution of sensory-driven activity to the dominant ongoing dynamics indexed by microstate C. This pattern is consistent with recent 7T fMRI findings in EC, which show a relative down-regulation of transmodal hubs alongside engagement of unimodal sensory systems (29). Together, these transition dynamics point to a redistribution of information flow in which sensory inputs are preferentially aligned with internally structured but self-minimal dynamics. More broadly, these findings are consistent with a hierarchical reconfiguration of precision weighting during EC, whereby sensory signals are preserved yet prevented from contributing to globally integrated, self-referential experiences.

The distribution of these effects across frequency sub-bands may provide further insight into the mechanisms and constraints underlying EC, although these observations should be interpreted cautiously given the small sample size. Converging evidence indicates that neural oscillations play a central role in coordinating large-scale communication, enabling the selective propagation and integration of information across distributed cortical networks through frequency-specific synchronization and dynamic gating mechanisms (58–60). More specifically, slower oscillations (e.g., delta, theta) are thought to support long-range coordination and top-down integration across distributed networks, whereas faster rhythms (e.g., beta, gamma) are more closely associated with local processing, feedforward signaling, and the propagation of sensory information (59, 61, 62). In our data, the reduction of microstate B was most pronounced in the theta and delta bands and was accompanied in delta by increased microstate A occurrence as well as elevated B-to-A transition probabilities. Together, this delta-theta pattern reinforces our interpretation of reduced coupling between sensory-driven and self-referential processing. From a computational perspective, it is consistent with a relative attenuation of top-down, self-related priors, alongside a shift toward sensory-driven updating, in line with the broader hierarchical reorganization of precision weighting inferred during EC (63).

By contrast, EC-related changes in microstate C were prominently expressed in the theta and beta, with beta additionally showing reduced microstate D parameters and fewer bidirectional D-B transitions. While microstate C may account for DMN/salience re-weighting activity, D is more generally linked to executive functions and sensorimotor integration (43). This theta–beta pattern therefore suggests a shift toward enhanced processing within local sensory-sensorimotor loops, accompanied by a reduced coupling between cognitive control and autobiographical systems during EC (64). Such a reconfiguration aligns with the functional segregation (29) and increased neural complexity (28) previously reported during EC. It further distinguishes EC from sleep or anaesthesia, which are typically characterized by a global shift toward slower rhythms and reduced complexity (35, 37, 65). In this context, the predominance of theta-band effects on duration of microstates B and C may reflect the engagement of sustained attentional control and long-range coordination across distributed networks (66, 67), supporting the extended routing and contextual modulation of information flow across hierarchical systems (68). This interpretation is further supported by microstate research suggesting that the temporal coordination of classes C and D may contribute to the balance between distributed cognitive systems (69, 70).

Finally, it is noteworthy that no specific microstate changes were identified in the alpha band. Classically, alpha-band oscillations are thought to reflect inhibitory and temporal gating processes that support core attentional functions such as suppression and selection, thereby enabling controlled knowledge access and semantic orientation (71). Within predictive processing accounts, alpha-band oscillations has been proposed to arise from hierarchical message passing under physiological conduction delays and to manifest as travelling waves (backward at rest) thereby indexing top-down modulation of sensory processing (63, 72). Interestingly, pharmacologic unconsciousness and sleep (N2) tend to “anteriorize” alpha power, that is, elevating alpha power in frontal regions of the brain while suppressing alpha power in occipital regions (73). This phenomenon has been mechanistically linked to GABAergic modulation under general anaesthesia, particularly in deep states such as propofol-induced unconsciousness, where cortical activity is strongly suppressed and frontal networks exhibit reduced functional engagement (73). In this context, anteriorization is thought to reflect the imposition of rhythmic activity onto a functionally downregulated cortex.

By contrast, EC is associated with a marked and widespread reduction in alpha power across cortical regions (7, 28), accompanied by a broad reduction in alpha-mediated coordination across large-scale networks (5), without clear evidence of a concomitant frontal alpha increase (28). From a mechanistic perspective, this divergence is consistent with the preservation of frontal lobe function, postural control, and volitional processes during EC, which stand in sharp contrast to the globally suppressed cortical state observed under anaesthesia. The absence of anteriorization in EC likely indicates that thalamo-frontal connectivity and top-down control mechanisms remain at least partially intact, allowing the thalamus to continue modulating higher-order cortical processes (74). Accordingly, the observed decrease in alpha power may reflect a distributed reconfiguration of alpha gating across cortical territories. In this view, alpha activity may be substantially downregulated at the macroscopic level while remaining functionally organized into narrower and more selective channels of coordination, consistent with the possibility that EC reflects an extreme form of large-scale gating operating across distributed networks.

Such a reconfiguration would not necessarily alter canonical scalp topographies and therefore may remain undetected at the level of alpha-specific microstate segmentation. Moreover, the absence of significant changes in the temporal parameters and transition probabilities of alpha-band microstates suggests that these modulations do not substantially reorganize the large-scale network alpha-dynamics but rather operate within existing repertoire of network states by shaping their internal excitability and cross-frequency information processing. This pattern is computationally consistent with a selective re-weighting of hierarchical functional influences, whereby alpha-mediated gating is relaxed at the global scale while becoming more selective and channel-specific, enabling flexible routing of information across distributed cortical systems without disrupting their underlying dynamical structure. Instead, EC would primarily operate by reconfiguring sensory sampling, neural computation and inter-area communication, thereby providing a plausible mechanistic route to the “reset” and post-EC wellbeing commonly reported by meditators (28, 75).

Beyond informing fundamental mechanisms of human consciousness, characterizing EC through fast EEG microstate dynamics may constitute a translational step for mental health, well-being, and human flourishing. While microstates track sub-second, whole-brain “atoms of thought” (30), the temporary absence of reportable experience during EC is followed by reproducible aftereffects, such as relief, clarity, and equanimity (9). These features are consistent with a trained and stabilized deployment of neural gating mechanisms enabling a controlled suspension of conscious integration. Although the present study was not designed to assess clinical outcomes, the identified sensor-level signatures may provide provisional targets for future advanced meditation-based interventions (76). Consistent with evidence linking meditation-related brain changes to improve emotional and cognitive function (77), time-frequency microstate analysis may offer a practical approach for monitoring meditative development (41) and its potential benefits for mental health and human flourishing (76).

Given the rarity of EC and the still-debated functional significance of EEG microstates, several limitations should be considered when interpreting these findings. Due to the limited number of practitioners worldwide who can reliably induce EC, we adopted an intensive dense-sampling design analogous to established small-N primate studies and human case studies (78). Accordingly, we estimated population-averaged effects using generalized estimating equations with an AR(1) correlation structure (79–81). Although our findings suggest that greater model generality (i.e., fewer, more global microstate classes) may better capture EC-related dynamics than increased specificity, the small sample (N = 5) likely contributed to identifying a limited number of microstates. Under these constraints, observed changes may reflect aggregates of collapsed sub-microstates with distinct functional roles. Notably, EC did not yield a novel, EC-specific microstate class. Instead, consistent with sleep and anesthesia research, EC appears to act on the large-scale integration of existing networks activities rather than their subdivision into finer-grained configurations (35–37). More broadly, links between microstates and specific underlying brain generators or functional systems remain tentative, as they rely on a limited number of source localization and EEG–fMRI studies. Accordingly, while canonical microstates are often related to large-scale networks (e.g., sensory, salience, or default-mode systems), such mappings remain indirect and should be considered as heuristic frameworks rather than definitive neurophysiological correspondences. Finally, although all participants completed a standardized EC phenomenology questionnaire, the absence of in-state self-reports limits inferences about subjective non-experience and its neural correlates. Accordingly, these findings should be regarded as preliminary and warrant replication in larger cohorts within the broader literature on advanced meditation and human consciousness (82).

Building on this initial characterization, future work will be essential to further delineate the neural mechanisms underlying EC. In particular, stimulus-evoked paradigms could assess the integrity and modulation of sensory processing during and immediately following EC. In clinical contexts such as akinetic mutism, auditory and somatosensory evoked potentials have demonstrated preserved early sensory responses despite impaired behavioral integration (83, 84), providing well-established benchmarks for dissociations between sensory encoding and conscious access. Modulation of auditory evoked potentials has also been reported in Vipassana meditation, consistent with changes in attentional deployment and sensory processing dynamics (85). Applying comparable paradigms to EC would allow direct tests of whether the inferred gating mechanisms operate downstream of intact early sensory responses, thereby bridging large-scale microstate dynamics with classical systems-level markers of consciousness. More broadly, multimodal approaches integrating EEG microstates with autonomic and interoceptive measures (such as cardiac dynamics or electrogastrogram activity), may provide a more synoptic understanding of how neural gating, sensory processing, and bodily regulation jointly contribute to the suspension and re-emergence of conscious experience during EC.

In conclusion, this study provides the first electrophysiological characterization of *extended cessation* of consciousness, revealing a distinctive scalp-level signature captured through microstate dynamics. EC appears as a specific form of phenomenological absence supported by a large-scale reorganization of functional brain dynamics, likely mediated by distributed alpha-dependent gating mechanisms that selectively constrain the integration of information into conscious experience. This reorganization was characterized by reduced self-related processing (microstate B), increased dominance of integrative default-mode/salience-related dynamics (microstate C), and altered routing of sensory information (microstate A). Frequency-specific analyses further refined this pattern, suggesting complementary contributions of slow and fast oscillatory dynamics to this large-scale reorganization. Together, these findings are consistent with a top-down re-weighting of precision within a predictive processing framework, whereby high-level self-related priors are attenuated while sensory signals remain active yet are prevented from being integrated into conscious experience. This account provides a parsimonious explanation for both the voluntary accessibility of EC and its characteristic afterglow. Beyond their theoretical implications, these findings identify time–frequency microstate dynamics as a promising electrophysiological window onto trained transitions in conscious experience and their potential relevance for mental health, well-being, and human flourishing.

## MATERIALS AND METHODS

### Participants

Participants were five advanced meditators (S1-S5, all male, mean age: 50 ± 12 years), each reporting being able to incline towards extended cessations as the target of their meditation practice. They had between 4 and 50 years (23 ± 16 years) of meditation experience with an estimated 9,000 to 25,000 lifetime practice hours. All were highly trained and capable of entering and exiting the EC state at predefined times, enabling experimental control over both onset and duration. Additional demographic and meditation-practice details are provided in the SI Appendix. The protocol and methodology used in this study are similar to that of previous published studies (29, 45). The protocol was approved by the Mass General Brigham Institutional Review Board, and all participants provided written informed consent.

### Experimental procedure

Participants completed approximately 45 minutes of EEG recordings across three EC sessions (except for Subject 1, SI Appendix). During each session, they attempted to maintain EC for approximately 15 minutes and indicated entry and exit using predefined behavioral markers when possible. Across sessions, EC durations ranged from 1 to 39 minutes (mean: 12.23 ± 8.15 min). Participants also completed a standardized phenomenology questionnaire designed to capture the typical phenomenology of their EC practice (Suppl. Fig. A). Two non-meditative control conditions (CC) were developed to compare against EC: (1) autobiographical memory retrieval and (2) structured mental counting. These tasks were designed to capture distinct forms of non-meditative cognitive engagement while minimizing the likelihood of spontaneous meditation (86), consistent with our previously published protocols (2, 3, 87). Two 8-min runs were collected for each control condition, providing 32 minutes of control condition data. High-density EEG was recorded under electrically shielded conditions using either a 96-channel actiCAP system (S1) or a 70/128-channel TRIUX system (S2-5), with sampling rates of 500 and 1000 Hz, respectively. Detailed procedures and acquisition parameters are provided in the SI Appendix.

### EEG Preprocessing

Offline EEG preprocessing was performed using EEGLab (88), a MATLAB-based toolbox. Processing included re-sampling to 500 Hz, band-pass filtering (1-50Hz), outlier detection and removal, independent component analysis (ICA), artefact rejection, interpolation of noisy channels and average re-referencing. Data were reduced to 64-channels according to the international standard 10–10 localization system. Each recording was FIR-filtered into canonical frequency bands: delta (1–3 Hz), theta (4–7 Hz), alpha (8–14 Hz), beta (15–30 Hz) and gamma (30-50 Hz). Finally, each EEG run was segmented into consecutive 1-min segments for statistical analyses, consistent with our previous work on advanced meditation (3, 29). Full preprocessing details are provided in the SI Appendix.

### Global Microstate Analysis

Microstate analysis was performed with freely available Cartool Software 3.70 (89). To identify the dominant scalp topographies across participants and conditions, we first performed a two-step microstate segmentation on broadband EEG data using hierarchical *k*-mean clusterings. Full clustering procedures and parameter settings are described in the SI Appendix. Clustering was applied to extract a common set of topographic maps (microstate classes) representing the most frequent spatial configurations across all datasets. The optimal number of clusters (k) was determined using a standard optimization metacriterion (52), yielding a set of global prototypical maps that best accounted for the spatial variance across participants and conditions. Considering the specific nature of the EC, we conducted additional segmentations at both the participant and condition levels to assess the robustness and specificity of the identified microstate classes. For each participant and each condition, independent k-means clustering was applied independently to the corresponding data subsets, with the optimization metacriterion used in each case to identify the optimal number of clusters. Spatial correlations were then computed by matching participant- and condition-specific centroids to the global microstate template, serving as a common reference, to quantify the consistency of topographic configurations across hierarchical levels of analysis. Meta-microstate analyses (46) were additionally performed to validate the correspondence between the identified templates and previously published microstate classes (Suppl. Table A). These group-level templates were then backfitted to the continuous broadband EEG data, allowing the extraction and comparison of temporal parameters across experimental conditions.

### EC-aligned Microstate Analysis

To further characterize frequency-specific effects associated with EC, we performed a second microstate analysis within canonical frequency bands (broadband, delta, theta, alpha, beta, gamma). Following established approaches (48), *k*-means clustering (*k* = 1:15, 200 repetitions, optimization meta-criterion) was applied within each frequency band to identify topographic maps that best explained spatio-temporal variance across participants and conditions. Given the observed dissimilarities between the global microstate template and EC-specific centroids (Suppl. Fig. B), the optimal number of clusters (k) was determined based on set of topographic maps that best accounted for the spatial variance in EC, while also considering GEV and clustering quality indices (mean criterion scores) across conditions. Accordingly, frequency-specific group-level maps were spatially correlated with the EC-derived centroids to identify prototypical topographies that best captured EC-specific effects while ensuring comparability across frequency bands. In addition, meta-microstate analyses were performed to validate the correspondence of the identified four-microstate templates with previously reported canonical microstate classes (Suppl. Table B) and to quantify the topographic relationship with the five-clustering solutions (Fig. 2).

Following this procedure, the four-microstate broadband-derived template was ultimately used as a common reference for backfitting across all frequency bands, given the high spatial correspondence observed between maps across frequency bands, conditions, and participants (Fig. 3). This approach ensured a consistent topographic framework for the extraction of temporal parameters, enabling direct comparison of frequency-specific dynamics while minimizing biases related to independent clustering solutions. As a control analysis, and in line with our a priori analysis plan, we additionally tested a frequency-specific segmentation approach in which band-limited maps were independently derived and fitted within each condition. This alternative approach yielded comparable results, confirming that the observed EC-related effects were robust to the choice of segmentation framework and were not driven by differences in clustering granularity. To specify the frequency domain, we labeled the microstate maps (MS) by adding the frequency band prefix (e.g., Bb-MS A, alpha-MS D, beta-MS C, etc.).

### Statistical Analysis

Statistical analyses were performed for microstate parameters and transitions probabilities using a custom Python pipeline based on statsmodels (v 0.14.4). Global explained variance (GEV, %), global field power (GFP, µV), mean duration (ms), occurrence frequency (s^-1^), time coverage (%), and transition probabilities (31) were separately analyzed using generalized estimating equations (GEE) with an autoregressive correlation structure (AR(1)) to account for repeated measurements within recording runs (79, 81). Wald χ^2^ tests assessed condition, microstate, and interaction effects (80). Condition was modeled with three categorial levels: EC, Counting, and Memory. We defined a combined control condition (CC) for a planned contrast, computed as the average of the Counting and Memory conditions, to test whether EC differed from the overall control-task context. Planned post-hoc contrasts were computed for both (i) the comparison of EC versus CC, and (ii) all pairwise comparisons between conditions (EC, Counting and Memory) to further characterize specific differences with control tasks and their functional relevance. To simplify results descriptions, statistics for (ii) are illustrated in SI Appendix (Suppl. Fig. C, D and Table C-H). False discovery rate (FDR) correction was applied using the Benjamini-Hochberg procedure (*q* = 0.05), correcting across microstates, conditions and frequency bands for each parameter. Effect sizes were reported in a model-appropriate manner. For logit-transformed parameters, effects are expressed as odds ratios (ORs), while for log-transformed parameters, effects are expressed as geometric mean ratios (GMRs). Additional modelling details are reported in the SI Appendix.

## Author Contributions

**D.A.E.Z.:** Data curation, Formal analysis, Software, Visualization, Writing – original draft. **W.F.Z.Y.:** Investigation, Formal analysis, Validation, Writing – review & editing. **A.R.:** Formal analysis, Software, Validation, Visualization, Writing – review & editing. **R.M.P.:** Investigation, Data curation, Project Administration, Writing – review & editing. **T.S.:** Methodology, Investigation, Resources, Writing – review & editing. **M.D.S.:** Conceptualization, Methodology, Investigation, Funding acquisition, Resources, Project Administration, Supervision, Writing – review & editing.

## Acknowledgments

We would like to express our gratitude to the meditation practitioners who agreed to take part in the study. We also acknowledge Avijit Chowdhury for helping with data collection and curation.

## Data and material availability

Data may be requested from the corresponding author and is subjected to the Massachusetts General Hospital Institutional Review Board’s guidelines and approval. All code used in this study may be made available by request from the corresponding author.

## References

1. A. K. Seth, T. Bayne, Theories of consciousness. Nat Rev Neurosci 23, 439–452 (2022).

2. A. Chowdhury, et al., Multimodal neurophenomenology of advanced concentration absorption meditation: An intensively sampled case study of Jhana. Neuroimage 305, 120973 (2025).

3. W. F. Z. Yang, et al., Intensive whole-brain 7T MRI case study of volitional control of brain activity in deep absorptive meditation states. Cereb Cortex 34, bhad408 (2024).

4. N. D. Schiff, F. Plum, The Role of Arousal and “Gating” Systems in the Neurology of Impaired Consciousness. Journal of Clinical Neurophysiology 17, 438 (2000).

5. R. van Lutterveld, A. Chowdhury, D. M. Ingram, M. D. Sacchet, Neurophenomenological Investigation of Mindfulness Meditation “Cessation” Experiences Using EEG Network Analysis in an Intensively Sampled Adept Meditator. Brain Topogr 37, 849–858 (2024).

6. A. Berkovich-Ohana, A case study of a meditation-induced altered state: increased overall gamma synchronization. Phenom Cogn Sci 16, 91–106 (2017).

7. A. Chowdhury, et al., Investigation of advanced mindfulness meditation “cessation” experiences using EEG spectral analysis in an intensively sampled case study. Neuropsychologia 190, 108694 (2023).

8. R. van Lutterveld, T. Cahaly, D. Ingram, M. D. Sacchet, Brain Criticality and Advanced Meditation “Cessation” Events: An Intensively Sampled Electroencephalography Case Study. [Preprint] (2025). Available at: https://papers.ssrn.com/abstract=5296987 [Accessed 30 January 2026].

9. R. E. Laukkonen, et al., Cessations of consciousness in meditation: Advancing a scientific understanding of nirodha samāpatti. Prog Brain Res 280, 61–87 (2023).

10. Buddhaghosa, The Path of Purification, Pariyatti Publishing (1999).

11. T. Sparby, M. D. Sacchet, The Third Wave of Meditation and Mindfulness Research and Implications for Challenging Experiences: Negative Effects, Transformative Psychological Growth, and Forms of Happiness. Mindfulness 16, 2156–2170 (2025).

12. M. D. Sacchet, M. Fava, E. L. Garland, Modulating self-referential processing through meditation and psychedelics: is scientific investigation of self-transcendence clinically relevant? World Psychiatry 23, 298–299 (2024).

13. L. Mudrik, et al., Unpacking the complexities of consciousness: Theories and reflections. Neuroscience & Biobehavioral Reviews 170, 106053 (2025).

14. T. Sparby, M. D. Sacchet, Toward a Scientific Model of Meditative Endpoints (MEND). Extremes of Human Experience and Development and the Case of Nirvana. [Preprint] (2025). Available at: https://osf.io/preprints/psyarxiv/ta3gn_v1/ [Accessed 24 June 2026].

15. M. H. Gunaratana, The Path of Serenity and Insight, Motilal Banarsidass (1985).

16. B. Nyanamoli Thera, B. Bodhi, The Middle Length Discourses of the Buddha: A Translation of the Majjhima Nikāya, Wisdom Publications (1995).

17. H. Noriaki, Nirodhasamapatti - Its Historical Meaning in the Vijñaptimatrata System. Indogaku Bukkyōgaku kenkyū 23, 1084–1074 (1975).

18. P. J. Griffths, On Being Mindless : Buddhist Meditation and the Mind-Body Problem (Bibliotheca Indo-Buddhica Series; No. 196), Open Court (1986).

19. D. Lusthaus, Buddhist Phenomenology: A Philosophical Investigation of Yogacara Buddhism and the Ch’eng Wei-shih Lun, Routledge (2002).

20. M. D. Sacchet, J. M. Lieberman, The neuroscience of advanced meditation: The promise of third wave meditation research. Neuron 114, 1545–1550 (2026).

21. J. Pujol, et al., Largest scale dissociation of brain activity at propofol-induced loss of consciousness. Sleep 44, zsaa152 (2021).

22. J. Pujol, et al., Mapping the neural systems driving breathing at the transition to unconsciousness. Neuroimage 246, 118779 (2022).

23. M. J. Wright, et al., Altered States of Consciousness are Prevalent and Insufficiently Supported Clinically: A Population Survey. Mindfulness 15, 1162–1175 (2024).

24. S. Prest, K. Berryman, Towards an Active Inference Account of Deep Meditative Deconstruction. [Preprint] (2024). Available at: https://osf.io/preprints/psyarxiv/d3gpf_v1/ [Accessed 19 May 2026].

25. H. Tal, M. Wright, S. Prest, L. Sandved-Smith, M. Sacchet, Active Inference, Computational Phenomenology, and Advanced Meditation: Toward the Formalization of the Experience of Meditation. [Preprint] (2025). Available at: https://osf.io/preprints/psyarxiv/pm5y2_v2/ [Accessed 27 December 2025].

26. N. Farb, et al., Interoception, contemplative practice, and health. Front. Psychol. 6 (2015).

27. A. K. Seth, K. J. Friston, Active interoceptive inference and the emotional brain. Philos Trans R Soc Lond B Biol Sci 371, 20160007 (2016).

28. K. Shinozuka, W. F. Z. Yang, R. M. Potash, T. Sparby, M. D. Sacchet, Neuroelectrophysiological correlates of extended cessation of consciousness in advanced meditators: A multimodal EEG and MEG study. [Preprint] (2025). Available at: https://www.biorxiv.org/content/10.1101/2025.09.19.677455v1 [Accessed 3 November 2025].

29. W. F. Z. Yang, et al., Endogenous suspension and reset of consciousness: 7T fMRI brain mapping of the extended cessation meditative endpoint. [Preprint] (2025). Available at: https://www.biorxiv.org/content/10.1101/2025.09.06.674021v2 [Accessed 3 November 2025].

30. D. Lehmann, H. Ozaki, I. Pal, EEG alpha map series: brain micro-states by space-oriented adaptive segmentation. Electroencephalogr Clin Neurophysiol 67, 271–288 (1987).

31. C. M. Michel, T. Koenig, EEG microstates as a tool for studying the temporal dynamics of whole-brain neuronal networks: A review. Neuroimage 180, 577–593 (2018).

32. E. Pipinis, et al., Association Between Resting-State Microstates and Ratings on the Amsterdam Resting-State Questionnaire. Brain Topogr 30, 245–248 (2017).

33. A. P. Zanesco, B. G. King, A. C. Skwara, C. D. Saron, Within and between-person correlates of the temporal dynamics of resting EEG microstates. Neuroimage 211, 116631 (2020).

34. A. P. Zanesco, E. Denkova, A. P. Jha, Self-reported Mind Wandering and Response Time Variability Differentiate Prestimulus Electroencephalogram Microstate Dynamics during a Sustained Attention Task. J Cogn Neurosci 33, 28–45 (2021).

35. V. Brodbeck, et al., EEG microstates of wakefulness and NREM sleep. Neuroimage 62, 2129–2139 (2012).

36. L. Bréchet, D. Brunet, L. Perogamvros, G. Tononi, C. M. Michel, EEG microstates of dreams. Sci Rep 10, 17069 (2020).

37. F. Artoni, et al., EEG microstate dynamics indicate a U-shaped path to propofol-induced loss of consciousness. Neuroimage 256, 119156 (2022).

38. R. Panda, et al., Temporal Dynamics of the Default Mode Network Characterize Meditation-Induced Alterations in Consciousness. Front Hum Neurosci 10, 372 (2016).

39. P. L. Faber, F. Travis, P. Milz, N. Parim, EEG microstates during different phases of Transcendental Meditation practice. Cogn Process 18, 307–314 (2017).

40. A. P. Zanesco, et al., Meditation training modulates brain electric microstates and felt states of awareness. Hum Brain Mapp 42, 3228–3252 (2021).

41. D. Zarka, et al., Electroencephalography microstates highlight specific mindfulness traits. Eur J Neurosci 59, 1753–1769 (2024).

42. A. Custo, et al., Electroencephalographic Resting-State Networks: Source Localization of Microstates. Brain Connect 7, 671–682 (2017).

43. P. Tarailis, T. Koenig, C. M. Michel, I. Griškova-Bulanova, The Functional Aspects of Resting EEG Microstates: A Systematic Review. Brain Topogr (2023). 10.1007/s10548-023-00958-9.

44. L. Bréchet, et al., Reconfiguration of Electroencephalography Microstate Networks after Breath-Focused, Digital Meditation Training. Brain Connect 11, 146–155 (2021).

45. D. Zarka, et al., Electroencephalography microstates highlight specific mindfulness traits. Eur J Neurosci 59, 1753–1769 (2024).

46. T. Koenig, et al., EEG-Meta-Microstates: Towards a More Objective Use of Resting-State EEG Microstate Findings Across Studies. Brain Topogr 37, 218–231 (2024).

47. T. Koenig, et al., Millisecond by millisecond, year by year: normative EEG microstates and developmental stages. Neuroimage 16, 41–48 (2002).

48. V. Férat, M. Seeber, C. M. Michel, T. Ros, Beyond broadband: Towards a spectral decomposition of electroencephalography microstates. Human Brain Mapping 43, 3047–3061 (2022).

49. M. T. Alkire, A. G. Hudetz, G. Tononi, Consciousness and anesthesia. Science 322, 876–880 (2008).

50. E. Altieri, L. Battaglini, Beyond Hyperexcitability: A Review of Neural Mechanisms in Charles Bonnet Syndrome. NeuroSci 7, 31 (2026).

51. C. P. Panayiotopoulos, Elementary visual hallucinations, blindness, and headache in idiopathic occipital epilepsy: differentiation from migraine. *Journal of Neurology*, Neurosurgery & Psychiatry 66, 536–540 (1999).

52. L. Bréchet, et al., Capturing the spatiotemporal dynamics of self-generated, task-initiated thoughts with EEG and fMRI. Neuroimage 194, 82–92 (2019).

53. F. Vellante, et al., Euthymic bipolar disorder patients and EEG microstates: a neural signature of their abnormal self experience? J Affect Disord 272, 326–334 (2020).

54. C. Ngo, et al., Microstate Dynamics of Focused Attention Meditation. Brain Topogr 39, 40 (2026).

55. A. P. Zanesco, E. Denkova, A. P. Jha, Self-reported Mind Wandering and Response Time Variability Differentiate Prestimulus Electroencephalogram Microstate Dynamics during a Sustained Attention Task. J Cogn Neurosci 33, 28–45 (2021).

56. L. Bréchet, C. M. Michel, EEG Microstates in Altered States of Consciousness. Front Psychol 13, 856697 (2022).

57. L. Bréchet, et al., Reconfiguration of Electroencephalography Microstate Networks after Breath-Focused, Digital Meditation Training. Brain Connect 11, 146–155 (2021).

58. S. Jahnke, R.-M. Memmesheimer, M. Timme, Oscillation-Induced Signal Transmission and Gating in Neural Circuits. PLoS Comput Biol 10, e1003940 (2014).

59. P. Fries, Rhythms for Cognition: Communication through Coherence. Neuron 88, 220–235 (2015).

60. S. van Bree, D. Levenstein, M. R. Krause, B. Voytek, R. Gao, Processes and measurements: a framework for understanding neural oscillations in field potentials. Trends in Cognitive Sciences 29, 448–466 (2025).

61. A. M. Bastos, et al., Visual areas exert feedforward and feedback influences through distinct frequency channels. Neuron 85, 390–401 (2015).

62. G. Michalareas, et al., Alpha-Beta and Gamma Rhythms Subserve Feedback and Feedforward Influences among Human Visual Cortical Areas. Neuron 89, 384–397 (2016).

63. K. J. Friston, Waves of prediction. PLOS Biology 17, e3000426 (2019).

64. A. Böttcher, et al., A dissociable functional relevance of theta- and beta-band activities during complex sensorimotor integration. Cereb Cortex 33, 9154–9164 (2023).

65. F. Von Wegner, et al., Complexity Measures for EEG Microstate Sequences: Concepts and Algorithms. Brain Topogr 37, 296–311 (2024).

66. M. S. Clayton, N. Yeung, R. Cohen Kadosh, The roles of cortical oscillations in sustained attention. Trends in Cognitive Sciences 19, 188–195 (2015).

67. J. Wei, Z. Zhang, Z. Yao, D. Ming, P. Zhou, Modulation of Sustained Attention by Theta-tACS over the Lateral and Medial Frontal Cortices. Neural Plast 2021, 5573471 (2021).

68. A. M. Bastos, M. Lundqvist, A. S. Waite, N. Kopell, E. K. Miller, Layer and rhythm specificity for predictive routing. Proceedings of the National Academy of Sciences 117, 31459–31469 (2020).

69. C. M. Michel, L. Bréchet, EEG microstates: from methodological foundations to clinical translation. Trends Neurosci 49, 433–449 (2026).

70. P. Tarailis, T. Koenig, C. M. Michel, I. Griškova-Bulanova, The Functional Aspects of Resting EEG Microstates: A Systematic Review. Brain Topogr 37, 181–217 (2024).

71. W. Klimesch, Alpha-band oscillations, attention, and controlled access to stored information. Trends Cogn Sci 16, 606–617 (2012).

72. A. Alamia, R. VanRullen, Alpha oscillations and traveling waves: Signatures of predictive coding? PLoS Biol 17, e3000487 (2019).

73. S. Vijayan, S. Ching, P. L. Purdon, E. N. Brown, N. J. Kopell, Thalamocortical mechanisms for the anteriorization of α rhythms during propofol-induced unconsciousness. J Neurosci 33, 11070–11075 (2013).

74. J. M. Shine, L. D. Lewis, D. D. Garrett, K. Hwang, The impact of the human thalamus on brain-wide information processing. Nat Rev Neurosci 24, 416–430 (2023).

75. A. Mierau, W. Klimesch, J. Lefebvre, State-dependent alpha peak frequency shifts: Experimental evidence, potential mechanisms and functional implications. Neuroscience 360, 146–154 (2017).

76. J. Galante, et al., A Framework for the Empirical Investigation of Mindfulness Meditative Development. Mindfulness 14, 1054–1067 (2023).

77. I. Sezer, D. A. Pizzagalli, M. D. Sacchet, Resting-state fMRI functional connectivity and mindfulness in clinical and non-clinical contexts: A review and synthesis. Neurosci Biobehav Rev 135, 104583 (2022).

78. A. Becker, The benefits of single-subject research designs and multi-methodological approaches for neuroscience research. Front. Hum. Neurosci. 17 (2023).

79. Y.-G. Wang, V. Carey, Working Correlation Structure Misspecification, Estimation and Covariate Design: Implications for Generalised Estimating Equations Performance. Biometrika 90, 29–41 (2003).

80. J. W. Hardin, J. M. Hilbe, Generalized Estimating Equations, 2nd Ed. (Chapman and Hall/CRC, 2012).

81. K.-Y. Liang, S. L. Zeger, Longitudinal data analysis using generalized linear models. Biometrika 73, 13–22 (1986).

82. R. S. Prakash, et al., Mindfulness Meditation and Network Neuroscience: Review, Synthesis, and Future Directions. Biol Psychiatry Cogn Neurosci Neuroimaging 10, 350–358 (2025).

83. B. Kotchoubey, M. Schneck, S. Lang, N. Birbaumer, Event-related brain potentials in a patient with akinetic mutism. Neurophysiologie Clinique/Clinical Neurophysiology 33, 23–30 (2003).

84. L. Naccache, et al., Preserved auditory cognitive ERPs in severe akinetic mutism: a case report. Brain Res Cogn Brain Res 19, 202–205 (2004).

85. B. R. Cahn, J. Polich, Meditation (Vipassana) and the P3a event-related brain potential. Int J Psychophysiol 72, 51–60 (2009).

86. Y.-Y. Tang, M. K. Rothbart, M. I. Posner, Neural correlates of establishing, maintaining, and switching brain states. Trends Cogn Sci 16, 330–337 (2012).

87. R. M. Potash, S. D. van Mil, M. Estarellas, A. Canales-Johnson, M. D. Sacchet, Integrated Phenomenology and Brain Connectivity Demonstrate Changes in Nonlinear Processing in Jhana Advanced Meditation. J Cogn Neurosci 37, 2260–2283 (2025).

88. A. Delorme, S. Makeig, EEGLAB: an open source toolbox for analysis of single-trial EEG dynamics including independent component analysis. J. Neurosci. Methods 134, 9–21 (2004).

89. D. Brunet, M. M. Murray, C. M. Michel, Spatiotemporal analysis of multichannel EEG: CARTOOL. Comput Intell Neurosci 2011, 813870 (2011).

