## Supplementary material for "Electrophysiological brain reconfiguration during meditation-induced extended cessation of consciousness: an EEG microstate study": Suppl.

### SUPPORTING INFORMATION APPENDIX

**SUPPLEMENTARY METHODS** ........................................................................ 1

**SUPPLEMENTARY FIGURES** ...................................................................... 7

**SUPPLEMENTARY TABLES** ......................................................................... 12

### SUPPLEMENTARY METHODS

#### Participants

Subject 1 (male, age: 52 years) is a long-term meditation practitioner with over 25 years of meditation experience and an estimated total practice amount at least of 20,000 hours at the time of data acquisition. Subject 2 (male, age: 32 years) had over 19 years of meditation experience with an estimated total practice amount of at least 15,000 hours. Subject 3 (male, age: 46 years) is an advanced meditator with over 4 years of meditation experience, and an estimated total practice amount of at least 9,000 hours. Subject 4 (male, age: 57 years) had over 50 years of meditation experience with total meditation experience of at least 10,000 hours. Subject 5 (male, age: 66 years) had over 19 years of meditation experience with total meditation experience at least of 25,000 hours. Lifetime practice hours are approximate self-reports based on duration and frequency of weekly practice and retreats. We also note that lifetime hours are not necessarily a direct measure of meditative expertise (1). The protocol and methodology used in this study are similar to that of previous published studies (2, 3). The Mass General Brigham IRB approved the study, and the participants provided informed consent.

#### Extended cessation

All participants were highly trained meditators capable of entering and exiting the EC state at predefined times, enabling experimental control over both onset and duration. During data collection, each participant completed approximately 45 minutes of EEG recordings across three EC sessions, except for Subject 1, who contributed a single EC run as part of a separate study. For each session, participants were instructed to meditate and volitionally enter EC for a target duration of 15 minutes. Given the absence of consciousness during EC, it was not possible for all participants to self-report their entrance; some indicated with an eye blink (S1), button press (S2, S4), or verbal report (S3, S5). Each run ended when the participant exited out of EC and indicated the exit with an eye blink (S1) or button press (S2, S3, S4, S5). Participants also completed a standardized EC phenomenology questionnaire designed to capture the typical phenomenology of their EC practice (Suppl. Fig. A). This questionnaire covered the following dimensions: preparation, onset, absence of consciousness, exit, and afterglow. The actual duration of each EC run lasted anywhere from 1 to 39 minutes. The mean duration of EC periods per subject was approximately 12.23 minutes (SD: 8.15) across sessions.

#### Non-meditative control conditions

Two non-meditative control conditions (CC) were developed to compare against EC: (1) a memory control condition in which participants were asked to reminisce the events of the past week and narrate them sub-vocally in their minds, without moving their lips and with their eyes closed, for eight minutes; and a (2) counting control condition, where the participants were asked to mentally count down in decrements of five from 10,000, without moving their lips and with their eyes closed, for 8 minutes. These two control conditions were used to capture distinct forms of non-meditative cognitive engagement (autobiographical recall vs. structured mental computation), thereby strengthening the specificity of comparisons with the experimental condition. These control condition were carefully designed to not induce meditative states and mirror our previously published protocols (4–6). As such, a resting-state control condition was not used, since experienced meditators may enter meditative states during this period (7). We collected two runs for each control condition, which provided 16 mins data for each control condition. However, for Subject 2, only a single run was performed for each control condition.

#### EEG data acquisition

Subjects' EEG data were acquired from two different facilities. S1's continuous EEG signals were recorded from a customized 96-channel actiCAP system using an actiCHamp amplifier (Brain Products GmbH, Gilching, Germany) inside an acoustically and electrically shielded booth for EEG data acquisition. EEG cap placement adhered to the standard 10-20 EEG positioning method. Impedances were kept below 5 kΩ. The ground (GND) channel was embedded in the cap and was located anterior and to the right of Channel 10, which roughly corresponds to electrode Fz. Channel 1 (Cz) served as the online reference channel during data acquisition. All signals were digitized at 500 Hz using BrainVision Recorder software (Brain Products). S2, S3, S4 and S5's continuous MEG-EEG signals were recorded on the TRIUX system at MGH with 70 (S2) and 128 EEG (S3, S4, S5) channels respectively. The MEG-EEG system was located in an Imedco magnetically shielded room, with a shielding factor of approximately 250,000 at 1Hz. The sampling frequency was 1000.0 Hz with a high bandpass of 0.1 Hz and a low bandpass of 330.0 Hz.

#### EEG Preprocessing

Offline preprocessing was performed using EEGLab (8), a MATLAB-based toolbox. Data were down sampled to 500 Hz and re-referenced to the average. They were filtered between 1 and 50 Hz using a finite-impulse response (FIR) filter (zero-phase, non-causal bandpass filter, -6 dB cutoff frequency, transition bandwidth: 1, transition length: 1651 samples). After visual inspection, outlying channels were rejected and noisy EEG segments were removed. Independent component analysis (ICA) was then performed using the infomax (runica) algorithm. Components reflecting ocular (blinks and saccades), muscular, and cardiac artefacts were classified using ICLabel (9) and rejected based on temporal, topographic features and frequency spectrum. Missing channels were interpolated using three-dimensional spherical splines (max. 10% of electrodes were interpolated for a single dataset). EEG data were then reduced to 64 electrodes according to the international standard 10–10 localization system. Each recording was FIR-filtered into traditional EEG frequency bands: delta (1–3 Hz, transition bandwidth: 1, transition length: 1651 samples), theta (4–7 Hz, transition bandwidth: 2, transition length: 827 samples), alpha (8–14 Hz, transition bandwidth: 2, transition length: 827 samples), beta (15–30 Hz, transition bandwidth: 3.75, transition length: 441 samples) and gamma (30-50 Hz, transition band width:7.5, transition length: 221 samples). Each EEG run was then segmented into one-minute segments for statistical analyses, similar to our previous work on advanced meditation (3, 5).

#### Microstate Segmentations

Microstate analysis was performed with freely available Cartool Software 3.70 (10). Data segmentation was applied across all frequency bands (broadband, delta, theta, alpha, beta and gamma), subjects (S1-5) and conditions (Memory, Counting, EC) following a two-step procedure. The first step consisted of computing independent clusters for each 1-min dataset (individual x condition x frequency band) using modified *k*-mean clustering. To this purpose, a subset of *k* (1:12) centroid map was randomly selected from the total set of voltage maps for each dataset, to use as initial centroids for clustering. Only voltage maps at time points of local maximum global field power (GFP) were considered for clustering to improve the signal-to-noise ratio. For each value of *k*, the *k* centroid maps were iteratively recomputed by averaging the voltage maps with which they showed the highest spatial correlation (minimum 0.5, the polarity of the maps was ignored). After 100 repetitions for each *k*-level (1:12), the *k* centroids with the maximum global explained variance (GEV) were selected. The optimal number of centroids among all possible values of *k* was determined using six independent optimization criteria (Gamma, Silhouettes, Davies and Bouldin, Point Biserial, Dunn and Krzanowski–Lai Index) merged into a single meta-criterion (11). At the end of this process, each dataset (individual x condition x frequency band) had a set of optimal clusters, each composed of *k* (1-12) centroids.

The second step consisted of conducting second-level *k*-means clustering (*k* = 1:15, 200 repetitions, optimization meta-criterion) on the centroids identified through clustering of the first-level voltage maps to identify the optimal global set of *k* common clusters that best explain the spatio-temporal variance of the EEG signals across participants and/or conditions within frequency bands (12). In each case, the common microstate maps obtained were then spatially back correlated (ignoring polarity) with the normalized map at each data point of the related original recordings. Thus, all samples of the original EEG were labelled with the microstate map with the highest correlation. EEG samples showing low spatial correlations (<0.5) with all microstate maps were left unassigned. To avoid artificially interrupting temporal segments of stable topography by noise, we used a sliding window (half size = 10; Besag factor = 10) (10). Finally, segments with a duration shorter than 10 samples (20 ms) were split into two parts and assigned to adjacent segments with the higher spatial correlation.

The spatio-temporal parameters of microstates were then calculated based on all time points with the same label. The GEV (%) is the percentage of the original EEG variance explained by each microstate map (weighted by the GFP at each moment in time). GFP reflects the maximal field strength and degree of synchronization among the neural generators contributing to the voltage maps for each microstate. The duration (ms) is the average time of continuous samples labelled to each microstate map. The occurrence ($s^{-1}$) is the frequency at which a given microstate appears in the continuous-time topographical series. The coverage (%) is the ratio of the time frames labelled to each microstate relative to the total time of the whole EEG recording (13). To study microstate syntax, we calculated a Markov matrix on the observed transitions between microstates and an expected Markov matrix by computing the expected transition probabilities given the distribution of microstate labels (13).

#### Meta-Microstate Analysis

To objectively validate the topographical configurations of the resulting global broadband templates, we performed a meta-microstate analysis using the MS-Template-Explorer tool (14). The broadband template maps from the current study were integrated with a database of 70 template sets from 58 published studies. All maps were interpolated to a common electrode montage using spherical splines and submitted to a modified k-means clustering algorithm (4 to 7 clusters) weighted by the number of subjects in each study to identify representative meta-microstate clusters. These meta-microstate templates were then back-fitted to both the database templates and the templates identified in our study. This back-fitting procedure was performed to quantitatively verify that our identified maps were centred within the established spatial variance (or "clouds") of previously published resting-state microstate types. The spatial relationship between our results and the global database was visualized using multidimensional scaling (MDS), where pairwise distances represent topographical dissimilarity derived from shared spatial variance (Fig. 2). We also performed spatial correlations (squared spatial correlation coefficients) between our templates and those of the 11 studies from the “Consciousness and its Disorders” research branch of the database (15–25) to assess shared spatial variance across all identified microstate classes within this branch of research (Suppl. Table A, B).

#### Statistical Analysis

Statistical analyses were performed separately for each microstate parameters (GEV, GFP, duration, occurrence, and coverage) and frequency band using a custom Python pipeline based on statsmodels (v 0.14.4). To account for the temporally correlated structure of the data arising from repeated measurements within recording runs, we used generalized estimating equations (GEEs) with a first-order autoregressive correlation structure (AR(1)). This GEE (AR(1)) structure yield population-averaged estimates while explicitly modeling the expected decay in correlation across adjacent 1-min segments within each subjects’ run, allowing reasonable inference while retaining all segmented observations (26, 27). To assess overall effects, we performed Wald $\chi^2$ tests on the fitted GEE models for the main effect of condition, the main effect of microstate, and their interaction (28). Condition was modeled with three categorial levels: EC, Counting, and Memory. We defined a combined control condition (CC) for a planned contrast, computed as the average of the Counting and Memory conditions, to test whether EC differed from the overall control-task context. Planned post-hoc contrasts were computed for both (i) the comparison of EC versus CC, and (ii) all pairwise comparisons between conditions (EC, Counting and Memory). This approach allowed us to assess the specificity of EC-related effects while retaining the separate EC–Counting, EC–Memory, and Counting–Memory contrasts, which were reported to further characterize differences between control tasks and their functional relevance. To simplify results descriptions, statistics for (ii) are illustrated in Supplementary (Suppl. Fig. C, D and Table C-H). False discovery rate (FDR) correction was applied using the Benjamini-Hochberg procedure ($q$ = 0.05), correcting across microstates, conditions and frequency bands for each parameter. Effect sizes were reported in a model-appropriate manner. For logit-transformed parameters, effects are expressed as odds ratios (ORs), while for log-transformed parameters, effects are expressed as geometric mean ratios (GMRs).

Microstate transition statistics were analyzed using the same GEE (AR(1)) framework as for the temporal microstate parameters, thereby ensuring consistent handling of temporal autocorrelation and repeated measurements. Transition values were defined as observed minus expected transition probabilities, where expected transitions were computed at random under the assumption of conserved microstate occurrences rates remain constant within each time segment (29). For within-condition analyses, observed-minus-expected transition values were modeled separately for each directed transition using Gaussian GEEs with an identity link, including an intercept only. For between-condition analyses, condition was again included as a three-level categorical predictor, allowing a planned EC vs. CC contrast (with CC defined as the average of Counting and Memory), as well as all pairwise contrasts between EC, Counting, and Memory (Suppl. Fig. C, E). In all cases, observations were clustered by subject-run, and an AR(1) working correlation structure was specified over temporally ordered segments within each subject–condition–run combination. Multiple-comparison correction across transitions was performed using FDR correction with the Benjamini–Hochberg procedure ($q$ = 0.05), applied separately for each contrast type and frequency band. Effect sizes are reported as mean differences in observed-minus-expected transition probabilities, or as differences of these quantities between conditions.

### REFERENCES


1. S. Ehmann, I. Sezer, I. N. Treves, J. D. E. Gabrieli, M. D. Sacchet, Mindfulness, cognition, and long-term meditators: Toward a science of advanced meditation. *Imaging Neuroscience* **3**, IMAG.a.82 (2025).
2. D. Zarka, *et al.*, Electroencephalography microstates highlight specific mindfulness traits. *Eur J Neurosci* **59**, 1753–1769 (2024).
3. W. F. Z. Yang, *et al.*, Endogenous suspension and reset of consciousness: 7T fMRI brain mapping of the extended cessation meditative endpoint. [Preprint] (2025). Available at: https://www.biorxiv.org/content/10.1101/2025.09.06.674021v2 [Accessed 3 November 2025].
4. R. M. Potash, S. D. van Mil, M. Estarellas, A. Canales-Johnson, M. D. Sacchet, Integrated Phenomenology and Brain Connectivity Demonstrate Changes in Nonlinear Processing in Jhana Advanced Meditation. *J Cogn Neurosci* **37**, 2260–2283 (2025).
5. W. F. Z. Yang, *et al.*, Intensive whole-brain 7T MRI case study of volitional control of brain activity in deep absorptive meditation states. *Cereb Cortex* **34**, bhad408 (2024).
6. A. Chowdhury, *et al.*, Multimodal neurophenomenology of advanced concentration absorption meditation: An intensively sampled case study of Jhana. *Neuroimage* **305**, 120973 (2025).
7. Y.-Y. Tang, M. K. Rothbart, M. I. Posner, Neural correlates of establishing, maintaining, and switching brain states. *Trends Cogn Sci* **16**, 330–337 (2012).
8. A. Delorme, S. Makeig, EEGLAB: an open source toolbox for analysis of single-trial EEG dynamics including independent component analysis. *J. Neurosci. Methods* **134**, 9–21 (2004).
9. L. Pion-Tonachini, K. Kreutz-Delgado, S. Makeig, ICLabel: An automated electroencephalographic independent component classifier, dataset, and website. *Neuroimage* **198**, 181–197 (2019).
10. D. Brunet, M. M. Murray, C. M. Michel, Spatiotemporal analysis of multichannel EEG: CARTOOL. *Comput Intell Neurosci* **2011**, 813870 (2011).
11. L. Bréchet, *et al.*, Capturing the spatiotemporal dynamics of self-generated, task-initiated thoughts with EEG and fMRI. *Neuroimage* **194**, 82–92 (2019).
12. V. Férat, M. Seeber, C. M. Michel, T. Ros, Beyond broadband: Towards a spectral decomposition of electroencephalography microstates. *Human Brain Mapping* **43**, 3047–3061 (2022).

13. C. M. Michel, T. Koenig, EEG microstates as a tool for studying the temporal dynamics of whole-brain neuronal networks: A review. *Neuroimage* **180**, 577–593 (2018).

14. T. Koenig, *et al.*, EEG-Meta-Microstates: Towards a More Objective Use of Resting-State EEG Microstate Findings Across Studies. *Brain Topogr* **37**, 218–231 (2024).

15. A. P. Zanesco, E. Denkova, A. P. Jha, Self-reported Mind Wandering and Response Time Variability Differentiate Prestimulus Electroencephalogram Microstate Dynamics during a Sustained Attention Task. *J Cogn Neurosci* **33**, 28–45 (2021).

16. L. Bréchet, D. Brunet, L. Perogamvros, G. Tononi, C. M. Michel, EEG microstates of dreams. *Sci Rep* **10**, 17069 (2020).

17. A. Custo, *et al.*, Electroencephalographic Resting-State Networks: Source Localization of Microstates. *Brain Connect* **7**, 671–682 (2017).

18. S. Denzer, S. Diezig, P. Achermann, F. W. Mast, T. Koenig, Electrophysiological (EEG) microstates during dream-like bizarre experiences in a naturalistic scenario using immersive virtual reality. *Eur J Neurosci* **60**, 5815–5830 (2024).

19. E. Toplutaş, F. Aydın, L. Hanoğlu, EEG Microstate Analysis in Patients with Disorders of Consciousness and Its Clinical Significance. *Brain Topogr* **37**, 377–387 (2024).

20. A. Pal, M. Behari, V. Goyal, R. Sharma, Study of EEG microstates in Parkinson's disease: a potential biomarker? *Cogn Neurodyn* **15**, 463–471 (2021).

21. P. Croce, A. Quercia, S. Costa, F. Zappasodi, EEG microstates associated with intra- and inter-subject alpha variability. *Sci Rep* **10**, 2469 (2020).

22. J. Britz, D. Van De Ville, C. M. Michel, BOLD correlates of EEG topography reveal rapid resting-state network dynamics. *Neuroimage* **52**, 1162–1170 (2010).

23. S. Diezig, S. Denzer, P. Achermann, F. W. Mast, T. Koenig, EEG Microstate Dynamics Associated with Dream-Like Experiences During the Transition to Sleep. *Brain Topogr* **37**, 343–355 (2024).

24. M. Murphy, R. Stickgold, M. E. Parr, C. Callahan, E. J. Wamsley, Recurrence of task-related electroencephalographic activity during post-training quiet rest and sleep. *Sci Rep* **8**, 5398 (2018).

25. J. N. Spring, E. F. Sallard, P. Trabucchi, G. P. Millet, J. Barral, Alterations in spontaneous electrical brain activity after an extreme mountain ultramarathon. *Biol Psychol* **171**, 108348 (2022).

26. K.-Y. Liang, S. L. Zeger, Longitudinal data analysis using generalized linear models. *Biometrika* **73**, 13–22 (1986).

27. Y.-G. Wang, V. Carey, Working Correlation Structure Misspecification, Estimation and Covariate Design: Implications for Generalised Estimating Equations Performance. *Biometrika* **90**, 29–41 (2003).

28. J. W. Hardin, J. M. Hilbe, *Generalized Estimating Equations*, 2nd Ed. (Chapman and Hall/CRC, 2012).

29. D. Lehmann, *et al.*, EEG microstate duration and syntax in acute, medication-naïve, first-episode schizophrenia: a multi-center study. *Psychiatry Research: Neuroimaging* **138**, 141–156 (2005).

**Related manuscript:** Electrophysiological brain reconfiguration during meditation-induced extended cessation of consciousness: an EEG microstate study

**FIGURES AND TABLES**

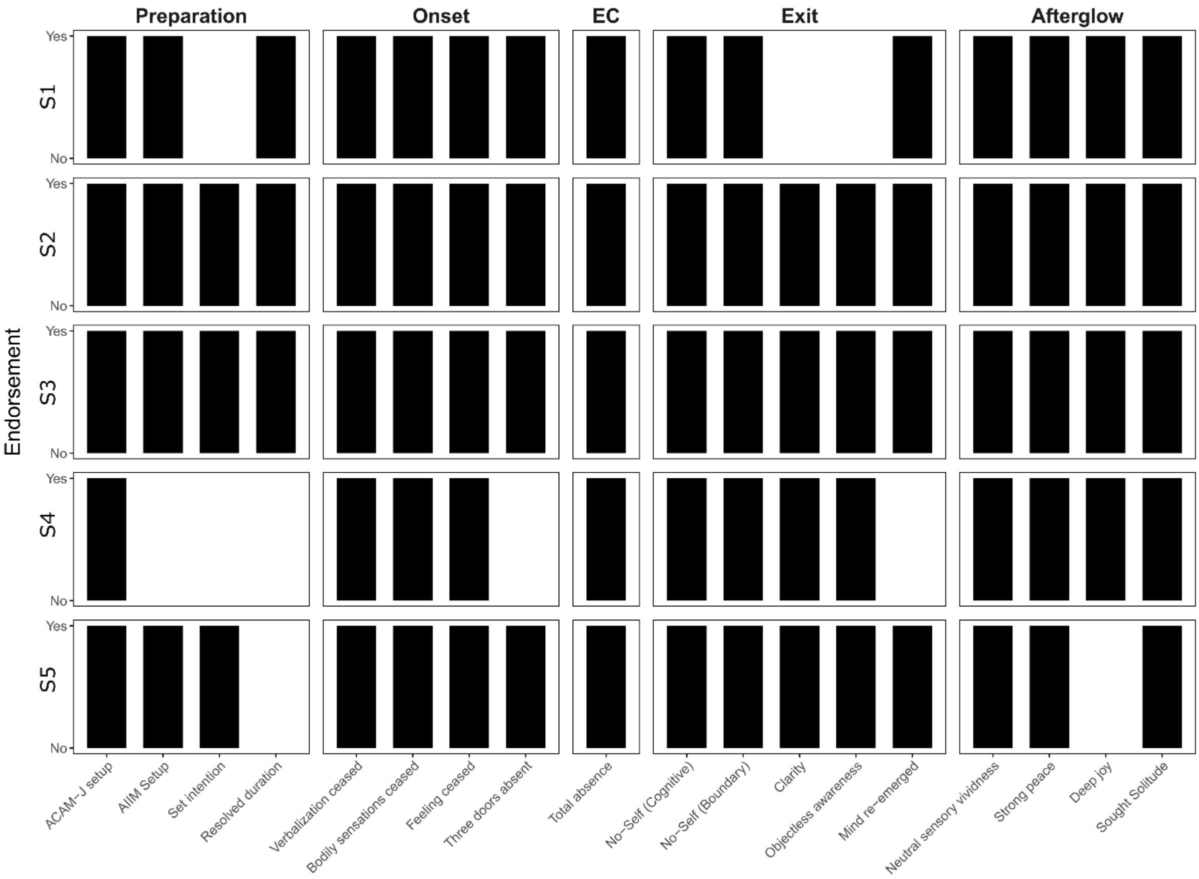


**Supplementary Figure A. Endorsement of EC phenomenology for each participant.** Participants all endorsed the phenomenological items of EC presented here, except Subject 4. Although previously trained to set up EC using ACAM-J, Subject 4 had since developed an alternative approach and no longer required ACAM-J for setup. As such, the phenomenology for various components throughout EC differ slightly for Subject 4. ACAM-J = Advanced concentration absorption meditation jhana, AIIM = advanced investigative insight meditation

**Related manuscript:** Electrophysiological brain reconfiguration during meditation-induced extended cessation of consciousness: an EEG microstate study

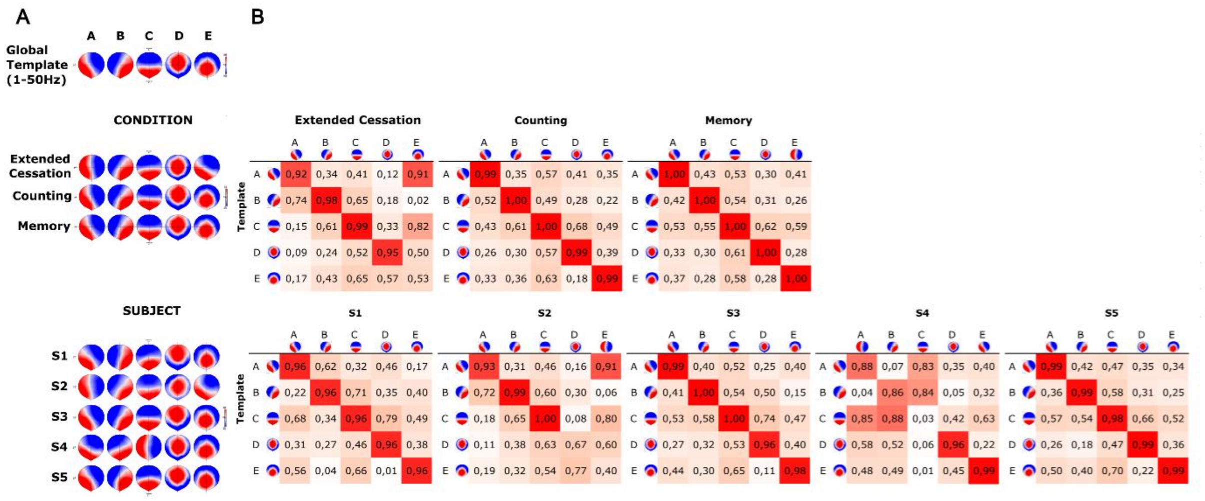


**Supplementary Figure B. Validation of the five-microstate solution across conditions and subjects.** (A) Canonical five-microstate solution (A–E) derived from the global broadband template (1–50 Hz), along with corresponding topographies obtained across experimental conditions (extended cessation, counting, and memory) and individual subjects (S1–S5). **(B)** Spatial correlation matrices between template maps and condition-specific, subject-specific, and frequency-specific maps. Diagonal values (highlighted in red) indicate high correspondence between matching configurations. The predominance of high diagonal correlations across matrices indicates a largely stable mapping of canonical microstates across conditions and participants. However, relatively elevated off-diagonal correlations, particularly between A-, C- and E-like configurations, indicate partial spatial overlap between these maps during EC and in individual participants (S2 and S4). Together, these results support the presence of five canonical microstate configurations across conditions, while highlighting increased spatial similarity between specific maps which may reduce their separability and favor the use of a more parsimonious four-microstate model to capture EC-related effects.

**Related manuscript:** Electrophysiological brain reconfiguration during meditation-induced extended cessation of consciousness: an EEG microstate study

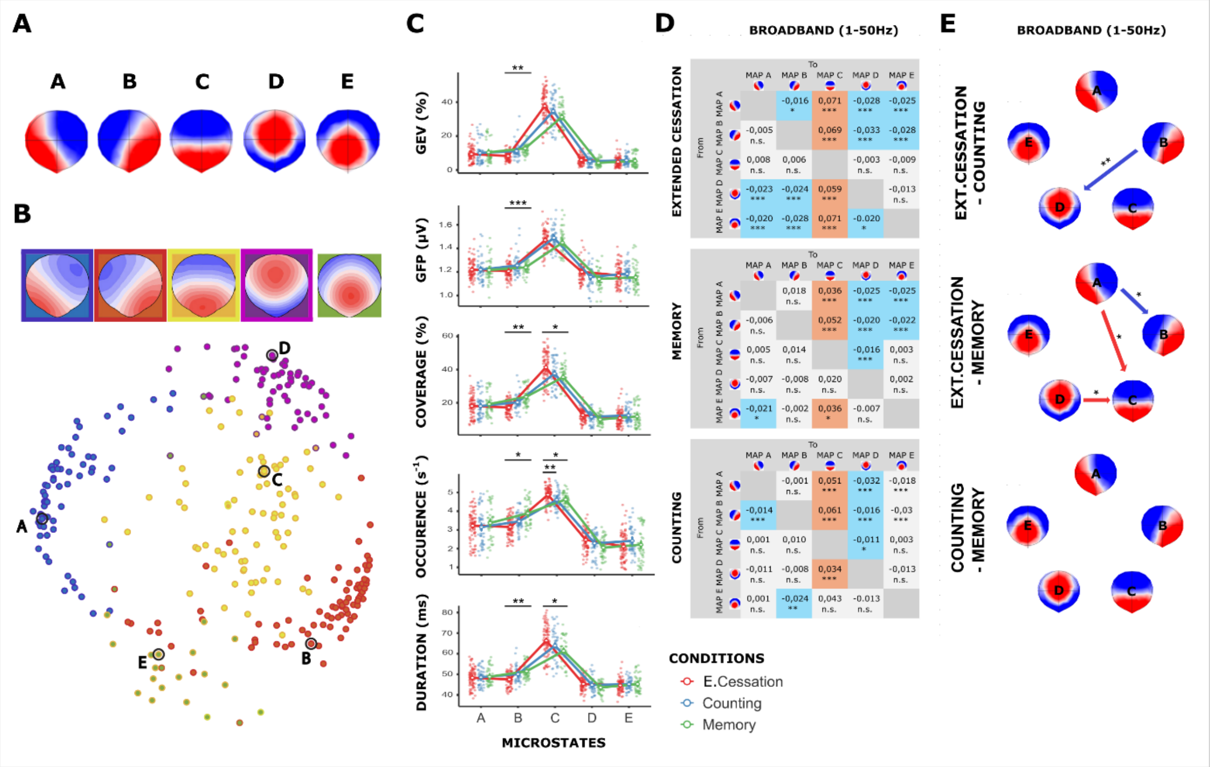


**Supplementary Figure C. Global broadband microstate organization and dynamics (five-microstate solution), separating the two control conditions.** **(A)** Group-level microstate topographies (A–E) identified using k-means clustering on broadband EEG data across participants and conditions. **(B)** Meta-microstate validation of the identified broadband templates in the four- and five-microstate solutions. **(C)** Microstate temporal parameters for the five-map solution, comparing Extended Cessation (red), Counting (blue) and Memory (green). **(D)** Markov transition probabilities between microstates for broadband data (1–50 Hz), shown separately for EC (top), Memory (middle) and Counting tasks (bottom). **(E)** Schematic representation of significant differences in transition probabilities between conditions (EC − counting; EC − memory; counting − memory). Arrows indicate direction and sign of modulation (red: increased; blue: decreased), summarizing the reorganization of microstate transitions in EC. Transitions involving microstate E were not significantly altered, indicating a limited contribution of this microstate to transition reorganization during EC.

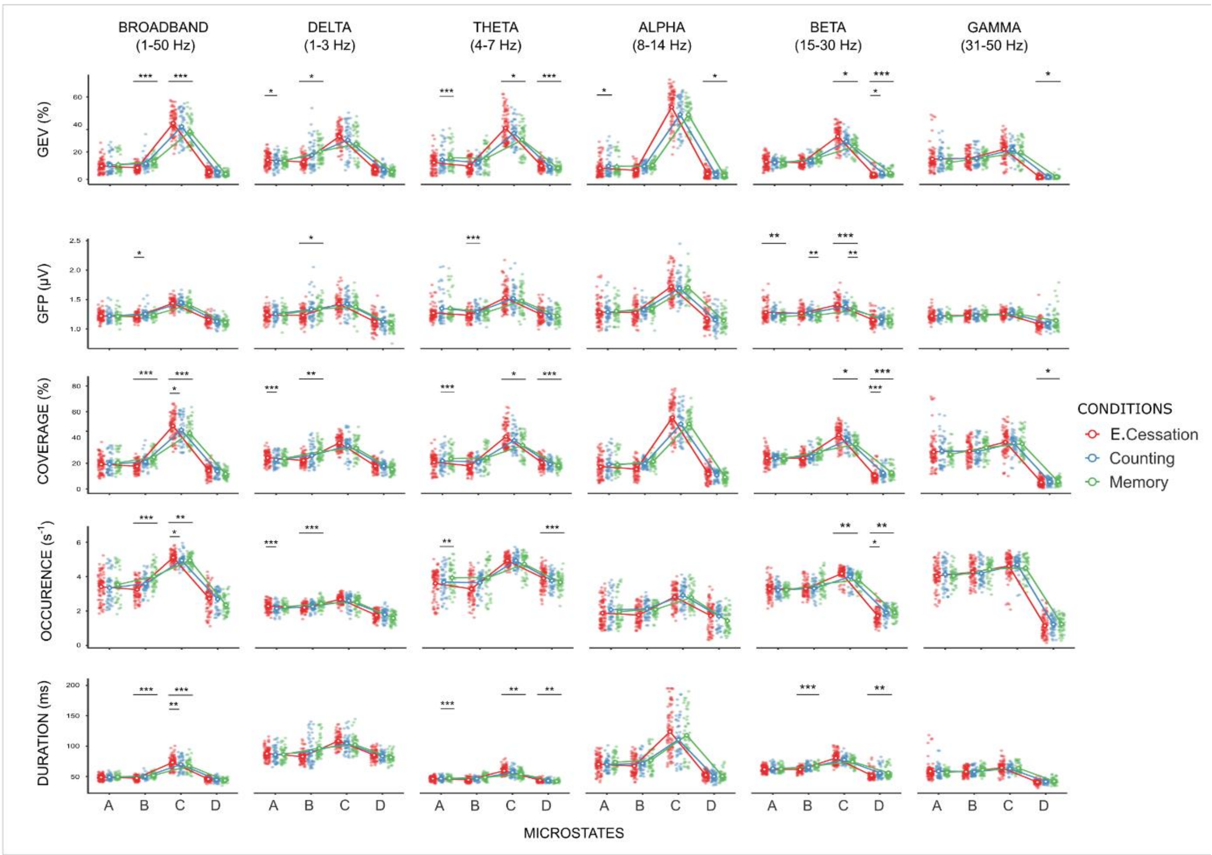


**Supplementary Figure D: Frequency-specific modulation of microstates parameters during extended cessation, compared to Counting and Memory tasks**. Canonical microstates characteristics are shown across broadband (1–50 Hz) and frequency-specific bands (delta, theta, alpha, beta, gamma), comparing extended cessation (red), Counting (blue) and Memory (green). Each row represents a distinct microstate parameter: global explained variance (GEV), global field power (GFP), time coverage, occurrence rate and mean duration. Statistics were performed using Wald $\chi^2$ tests from generalized estimating equation (GEE) models with autoregressive correlation structure of order 1 (AR(1)). Asterisks indicate FDR-corrected significance level (* $p < 0.05$, ** $p < 0.01$, *** $p < 0.001$).

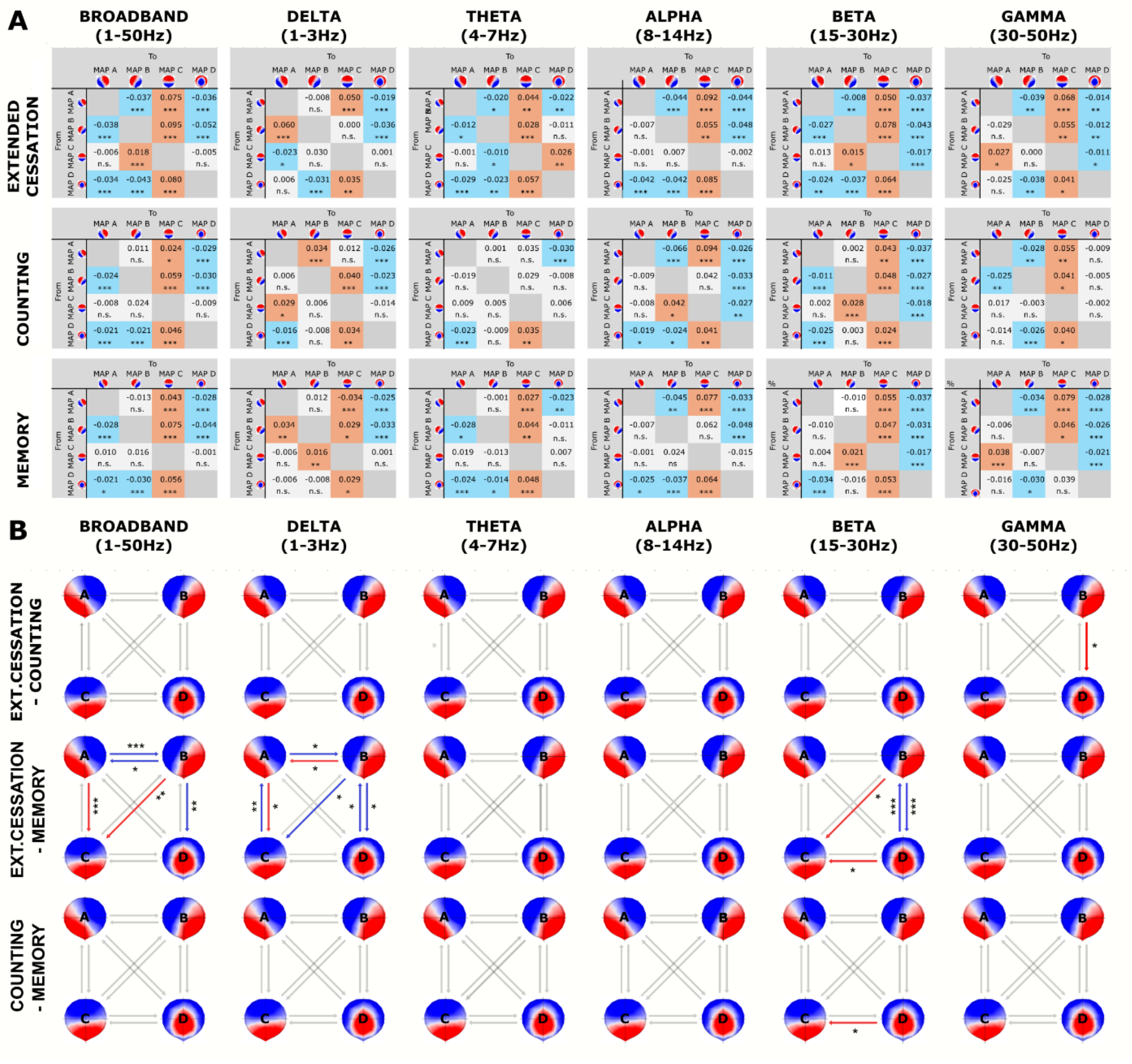


**Supplementary Figure E: Reconfiguration of microstate transition dynamics during Extended Cessation (EC), compared to Counting and Memory tasks. (A)** Transition probabilities differences (observed-minus-expected) within each condition across broadband (1-50Hz), delta (1-3Hz), theta (4-7Hz), alpha (8-14Hz), Beta (15-30Hz) and Gamma (30-50Hz) frequency bands. Each matrix represents transition likelihoods between canonical microstates (rows: "from", columns: "to"), with color-coded values. Positive values (red) indicate more frequent-than-random transitions, and negative values (blue) indicate less frequent-than-random transitions. Color and text indicate statistically significant differences as calculated by FDR-adjusted tests. Asterisks indicate significance level (* $p < 0.05$, ** $p < 0.01$, *** $p < 0.001$). **(B)** Network graphs of the between-condition comparisons (EC − counting; EC − memory; counting − memory) of the transition probabilities across the four canonical microstates for each frequency bands. Arrow indicates transition that are significantly increased (red) or decreased (blue), as calculated by FDR-adjusted Wald $\chi^2$ tests.

Table A: Similarity scores for the five-microstate solution (% shared variance; squared spatial correlation) compared to templates across studies in the Consciousness and its Disorders category.

| Study | MS A | MS B | MS C | MS D | MS E |
|---|---|---|---|---|---|
| Artoni (2022) | 92,5 | 92,5 | 93,5 | 87,1 | 92,9 |
| Bréchet (2020) | 72,1 | 67,6 | 87,9 (*E*) | 71,1 (*C*) | 75,4 (*D*) |
| Britz (2010) | 91,7 | 90,1 | 95,6 | 73,4 | - |
| Croce (2020) | 93,8 | 41,2 | 91,2 (*B*) | 67,8 (*C*) | 93,3 (*D*) |
| Custo (2017) | 92,9 | 88,1 | 98,4 | 94,5 | 84,3 (*F*) |
| Denzer (2024) | 58,4 | 44,4 | 87,6 (*B*) | 71,0 | 75,6 (*C*) |
| Diezig (2022) | 92,4 | 96,2 | 95,8 | 82,3 (*E*) | 88,7 (*D*) |
| Murphy (2018) | 81,2 (*F*) | 92,0 | 97,8 | 86,8 | 87,7 (*E*) |
| Pal (2021) | 48,9 (*E*) | 88,5 (*F*) | 93,3 (*B*) | 76,9 (*A*) | 71,8 (*C*) |
| Spring (2022) | 93,5 | 93,9 | 97,1 | 89,9 | - |
| Toplutas (2023) | 85,1 | 90,3 | 82,0 | 77,0 | - |
| Zanesco (2021) | 96,6 | 93,5 | 81,2 | 81,4 | 71,1 |
| **Average** | **83,26** | **81,5** | **91,8** | **79,9** | **82,3** |

*Note.* Each cell reports the best-matching template in the referenced study against that of the current study; parenthetical letters indicate the referenced study's label when it differs from the current labeling. For Toplutas (2023), values reflect the mean of three available template sets.

Table B: Similarity scores for the four-microstate solution (% shared variance; squared spatial correlation) compared to templates across studies in the Consciousness and its Disorders category.

| Study | MS A | MS B | MS C | MS D |
|---|---|---|---|---|
| Artoni (2022) | 93,7 | 91,9 | 94,0 | 82,8 |
| Bréchet (2020) | 75,4 | 67,7 | 92,5 (*E*) | 66,5 |
| Britz (2010) | 92,3 | 89,2 | 95,7 | 69,3 |
| Croce (2020) | 91,2 | 43,4 | 94,3 (*B*) | 46,1 (*C*) |
| Custo (2017) | 94,1 | 88,4 | 96,5 | 94,3 |
| Denzer (2024) | 58,6 | 44,1 | 82,7 (*B*) | 85 |
| Diezig (2022) | 93,1 | 95,1 | 97,9 | 63,9 (*E*) |
| Murphy (2018) | 81,2 (*E*) | 92,9 | 98,3 | 73,3 |
| Pal (2021) | 52,6 (*E*) | 89,2 (*F*) | 91,3 (*B*) | 56,8 (*A*) |
| Spring (2022) | 95,5 | 92,7 | 97,8 | 96,6 |
| Toplutas (2023) | 84,1 | 89,9 | 86,4 | 69,3 |
| Zanesco (2021) | 97,1 | 93 | 86,6 | 63,4 (*E*) |
| **Average** | **84,1** | **81,4** | **92,8** | **72,3** |

*Note.* Each cell reports the best-matching template in the referenced study against that of the current study; parenthetical letters indicate the referenced study's label when it differs from the current labeling. For Toplutas (2023), values reflect the mean of three available template sets.

Table C: Wald $\chi^2$ omnibus tests from GEE models (AR(1)) for the five-microstate solution in Broadband. $p$ values are uncorrected; $q$ values are FDR-corrected.

| Band | Effect | $\chi^2$ | df | $p$ | $q$ (FDR) |
|---|---|---|---|---|---|
| | Condition | 0.74 | 2 | 0.692 | 0.948 |
| GEV | Microstate | 69,902.78 | 3 | $< 0.001$ | $< 0.001$ |
| | Condition × Microstate | 195.56 | 6 | $< 0.001$ | $< 0.001$ |
| | Condition | 0.11 | 2 | 0.948 | 0.948 |
| GFP | Microstate | 3,183.02 | 3 | $< 0.001$ | $< 0.001$ |
| | Condition × Microstate | 4.55 | 6 | 0.804 | 0.804 |
| | Condition | 1.81 | 2 | 0.404 | 0.948 |
| Coverage | Microstate | 7,546.53 | 3 | $< 0.001$ | $< 0.001$ |
| | Condition × Microstate | 154.71 | 6 | $< 0.001$ | $< 0.001$ |
| | Condition | 2.94 | 2 | 0.229 | 0.948 |
| Occurrence | Microstate | 225.76 | 3 | $< 0.001$ | $< 0.001$ |
| | Condition × Microstate | 96.68 | 6 | $< 0.001$ | $< 0.001$ |
| | Condition | 0.14 | 2 | 0.932 | 0.948 |
| Duration | Microstate | 26,949.46 | 3 | $< 0.001$ | $< 0.001$ |
| | Condition × Microstate | 15.89 | 6 | 0.044 | 0.055 |

*Note.* $q$ values are Benjamini–Hochberg FDR-corrected for each effect type (Condition, Microstate, Interaction) across microstate parameters.

Table D: Wald $\chi^2$ omnibus tests from GEE models (AR(1)) for the GEV of the four-microstate solution across frequency bands. $p$ values are uncorrected; $q$ values are FDR-corrected.

| Band | Effect | $\chi^2$ | df | $p$ | $q$ (FDR) |
|---|---|---|---|---|---|
| Broadband | Condition | 1.70 | 2 | 0.427 | 0.512 |
| | Microstate | 232.90 | 3 | $< 0.001$ | $< 0.001$ |
| | Condition × Microstate | 366.12 | 6 | $< 0.001$ | $< 0.001$ |
| Delta | Condition | 44.22 | 2 | $< 0.001$ | $< 0.001$ |
| | Microstate | 1,129.35 | 3 | $< 0.001$ | $< 0.001$ |
| | Condition × Microstate | 290.57 | 6 | $< 0.001$ | $< 0.001$ |
| Theta | Condition | 35.42 | 2 | $< 0.001$ | $< 0.001$ |
| | Microstate | 2,112.25 | 3 | $< 0.001$ | $< 0.001$ |
| | Condition × Microstate | 208.54 | 6 | $< 0.001$ | $< 0.001$ |
| Alpha | Condition | 7.08 | 2 | 0.029 | 0.058 |
| | Microstate | 652.63 | 3 | $< 0.001$ | $< 0.001$ |
| | Condition × Microstate | 22.31 | 6 | 0.001 | 0.002 |
| Beta | Condition | 3.15 | 2 | 0.208 | 0.311 |
| | Microstate | 52.70 | 3 | $< 0.001$ | $< 0.001$ |
| | Condition × Microstate | $> 10^{15}$ | 6 | $< 0.001$ | $< 0.001$ |
| Gamma | Condition | 0.70 | 2 | 0.703 | 0.703 |
| | Microstate | 157.27 | 3 | $< 0.001$ | $< 0.001$ |
| | Condition × Microstate | 7.07 | 6 | 0.315 | 0.378 |

*Note.* $q$ values are Benjamini–Hochberg FDR-corrected within parameter for each effect type (Condition, Microstate, Interaction) across frequency bands.

Table E: Wald $\chi^2$ omnibus tests from GEE models (AR(1)) for the GFP of the four-microstate solution across frequency bands. $p$ values are uncorrected; $q$ values are FDR-corrected.

| Band | Effect | $\chi^2$ | df | $p$ | $q$ (FDR) |
|---|---|---|---|---|---|
| Broadband | Condition | 0.06 | 2 | 0.972 | 0.972 |
| | Microstate | 867.68 | 3 | $< 0.001$ | $< 0.001$ |
| | Condition × Microstate | 7.69 | 6 | 0.261 | 0.261 |
| Delta | Condition | 2.04 | 2 | 0.361 | 0.433 |
| | Microstate | 295.14 | 3 | $< 0.001$ | $< 0.001$ |
| | Condition × Microstate | $8.14 \times 10^{14}$ | 6 | $< 0.001$ | $< 0.001$ |
| Theta | Condition | 5.17 | 2 | 0.075 | 0.151 |
| | Microstate | 2,030.88 | 3 | $< 0.001$ | $< 0.001$ |
| | Condition × Microstate | 17,050.14 | 6 | $< 0.001$ | $< 0.001$ |
| Alpha | Condition | 3.47 | 2 | 0.177 | 0.265 |
| | Microstate | 2,361.32 | 3 | $< 0.001$ | $< 0.001$ |
| | Condition × Microstate | 12.17 | 6 | 0.058 | 0.070 |
| Beta | Condition | 52.41 | 2 | $< 0.001$ | $< 0.001$ |
| | Microstate | 1,563.35 | 3 | $< 0.001$ | $< 0.001$ |
| | Condition × Microstate | $8.27 \times 10^{12}$ | 6 | $< 0.001$ | $< 0.001$ |
| Gamma | Condition | 8.89 | 2 | 0.012 | 0.035 |
| | Microstate | 272.75 | 3 | $< 0.001$ | $< 0.001$ |
| | Condition × Microstate | 43.13 | 6 | $< 0.001$ | $< 0.001$ |

*Note.* $q$ values are Benjamini–Hochberg FDR-corrected within parameter for each effect type (Condition, Microstate, Interaction) across frequency bands.

Table F: Wald $\chi^2$ omnibus tests from GEE models (AR(1)) for the coverage of the four-microstate solution across frequency bands. $p$ uncorrected; $q$ FDR-corrected.

| Band | Effect | $\chi^2$ | df | $p$ | $q$ (FDR) |
|---|---|---|---|---|---|
| | Condition | 2.81 | 2 | 0.245 | 0.295 |
| Broadband | Microstate | 174.50 | 3 | $< 0.001$ | $< 0.001$ |
| | Condition $\times$ Microstate | 117.04 | 6 | $< 0.001$ | $< 0.001$ |
| | Condition | 35.63 | 2 | $< 0.001$ | $< 0.001$ |
| Delta | Microstate | 1,374.37 | 3 | $< 0.001$ | $< 0.001$ |
| | Condition $\times$ Microstate | 301.39 | 6 | $< 0.001$ | $< 0.001$ |
| | Condition | 17.80 | 2 | $< 0.001$ | $< 0.001$ |
| Theta | Microstate | 894.24 | 3 | $< 0.001$ | $< 0.001$ |
| | Condition $\times$ Microstate | 794.30 | 6 | $< 0.001$ | $< 0.001$ |
| | Condition | 4.60 | 2 | 0.100 | 0.150 |
| Alpha | Microstate | 435.52 | 3 | $< 0.001$ | $< 0.001$ |
| | Condition $\times$ Microstate | 6.58 | 6 | 0.361 | 0.361 |
| | Condition | 4.82 | 2 | 0.090 | 0.150 |
| Beta | Microstate | 54.37 | 3 | $< 0.001$ | $< 0.001$ |
| | Condition $\times$ Microstate | $8.11 \times 10^{13}$ | 6 | $< 0.001$ | $< 0.001$ |
| | Condition | 0.89 | 2 | 0.640 | 0.640 |
| Gamma | Microstate | 205.88 | 3 | $< 0.001$ | $< 0.001$ |
| | Condition $\times$ Microstate | 8.65 | 6 | 0.194 | 0.233 |

*Note.* $q$ values are Benjamini–Hochberg FDR-corrected within parameter for each effect type (Condition, Microstate, Interaction) across frequency bands.

Table G: Wald $\chi^2$ omnibus tests from GEE models (AR(1)) for the occurrence of the four-microstate solution across frequency bands. $p$ values are uncorrected; $q$ values are FDR-corrected.

| Band | Effect | $\chi^2$ | df | $p$ | $q$ (FDR) |
|---|---|---|---|---|---|
| Broadband | Condition | 2.48 | 2 | 0.290 | 0.382 |
| | Microstate | 370.01 | 3 | $< 0.001$ | $< 0.001$ |
| | Condition × Microstate | 45.99 | 6 | $< 0.001$ | $< 0.001$ |
| Delta | Condition | 38.65 | 2 | $< 0.001$ | $< 0.001$ |
| | Microstate | 1,094.40 | 3 | $< 0.001$ | $< 0.001$ |
| | Condition × Microstate | 154.20 | 6 | $< 0.001$ | $< 0.001$ |
| Theta | Condition | 9.30 | 2 | 0.010 | 0.029 |
| | Microstate | 426.28 | 3 | $< 0.001$ | $< 0.001$ |
| | Condition × Microstate | 22.18 | 6 | 0.001 | 0.001 |
| Alpha | Condition | 2.29 | 2 | 0.319 | 0.382 |
| | Microstate | 225.76 | 3 | $< 0.001$ | $< 0.001$ |
| | Condition × Microstate | 8.39 | 6 | 0.211 | 0.211 |
| Beta | Condition | 4.35 | 2 | 0.114 | 0.227 |
| | Microstate | 112.99 | 3 | $< 0.001$ | $< 0.001$ |
| | Condition × Microstate | $4.36 \times 10^{13}$ | 6 | $< 0.001$ | $< 0.001$ |
| Gamma | Condition | 0.89 | 2 | 0.642 | 0.642 |
| | Microstate | 353.83 | 3 | $< 0.001$ | $< 0.001$ |
| | Condition × Microstate | 26.22 | 6 | $< 0.001$ | $< 0.001$ |

*Note.* $q$ values are Benjamini–Hochberg FDR-corrected within parameter for each effect type (Condition, Microstate, Interaction) across frequency bands.

Table H: Wald $\chi^2$ omnibus tests from GEE models (AR(1)) for the duration of the four-microstate solution across frequency bands. $p$ values are uncorrected; $q$ values are FDR-corrected.

| Band | Effect | $\chi^2$ | df | $p$ | $q$ (FDR) |
|---|---|---|---|---|---|
| Broadband | Condition | 6.57 | 2 | 0.037 | 0.085 |
| | Microstate | 131.71 | 3 | $< 0.001$ | $< 0.001$ |
| | Condition × Microstate | 42.74 | 6 | $< 0.001$ | $< 0.001$ |
| Delta | Condition | 6.32 | 2 | 0.042 | 0.085 |
| | Microstate | 422.49 | 3 | $< 0.001$ | $< 0.001$ |
| | Condition × Microstate | $> 10^{15}$ | 6 | $< 0.001$ | $< 0.001$ |
| Theta | Condition | 143.33 | 2 | $< 0.001$ | $< 0.001$ |
| | Microstate | 2,317.09 | 3 | $< 0.001$ | $< 0.001$ |
| | Condition × Microstate | $2.18 \times 10^{13}$ | 6 | $< 0.001$ | $< 0.001$ |
| Alpha | Condition | 2.82 | 2 | 0.244 | 0.366 |
| | Microstate | 470.33 | 3 | $< 0.001$ | $< 0.001$ |
| | Condition × Microstate | 23.75 | 6 | $< 0.001$ | $< 0.001$ |
| Beta | Condition | 0.58 | 2 | 0.747 | 0.747 |
| | Microstate | 64.27 | 3 | $< 0.001$ | $< 0.001$ |
| | Condition × Microstate | $3.62 \times 10^{12}$ | 6 | $< 0.001$ | $< 0.001$ |
| Gamma | Condition | 1.06 | 2 | 0.590 | 0.708 |
| | Microstate | 173.42 | 3 | $< 0.001$ | $< 0.001$ |
| | Condition × Microstate | 147.29 | 6 | $< 0.001$ | $< 0.001$ |

*Note.* $q$ values are Benjamini–Hochberg FDR-corrected within parameter for each effect type (Condition, Microstate, Interaction) across frequency bands.
